# *rox*: A statistical model for regression with missing values

**DOI:** 10.1101/2022.04.15.488427

**Authors:** Mustafa Buyukozkan, Elisa Benedetti, Jan Krumsiek

## Abstract

High-dimensional omics datasets frequently contain missing data points, which typically occur due to concentrations below the limit of detection (LOD) of the profiling platform. The presence of such missing values significantly limits downstream statistical analysis and result interpretation. Two common techniques to deal with this issue include the removal of samples with missing values, and imputation approaches which substitute the missing measurements with reasonable estimates. Both approaches, however, suffer from various shortcomings and pitfalls. In this paper, we present “*rox*”, a novel statistical model for the analysis of omics data with missing values without the need for imputation. The model directly incorporates missing values as “low” concentrations into the calculation. We show the superiority of *rox* over common approaches on simulated data and on six metabolomics datasets. Fully leveraging the information contained in LOD-based missing values, *rox* provides a powerful tool for the statistical analysis of omics data.

## 1 Introduction

High-dimensional molecular datasets, such as metabolomics, proteomics, glycomics and microbiomics, typically contain a substantial amount of “missing values”, that is, measurement points for which the experimental platform did not return any quantified value [1, 2, 3]. Any analysis workflow applied to data with missing values needs to deal with this issue, since most common statistical approaches do not allow for the absence of data points. Missing values in omics data usually occur due to abundances below the instrument sensitivity, the so-called limit of detection (LOD) [3] (Figure 1A). In addition to the obvious loss of information, the presence of missing values interferes with distributional assumptions for statistical analysis. For example, metabolomics measurements are generally log-normally distributed [4], and therefore LOD-based missing values will obfuscate the lower tail of the distribution. In microbiome data, which are compositional in nature [5], left-truncation will lead to an artificial overrepresentation of the most abundant species. Further complicating the issue, we have previously shown that LOD effects are not always strict and can occur in blurry fashion, where lower concentration values increase the chance of a value being reported as missing rather than depending on a strict threshold [3].

**Figure 1:**
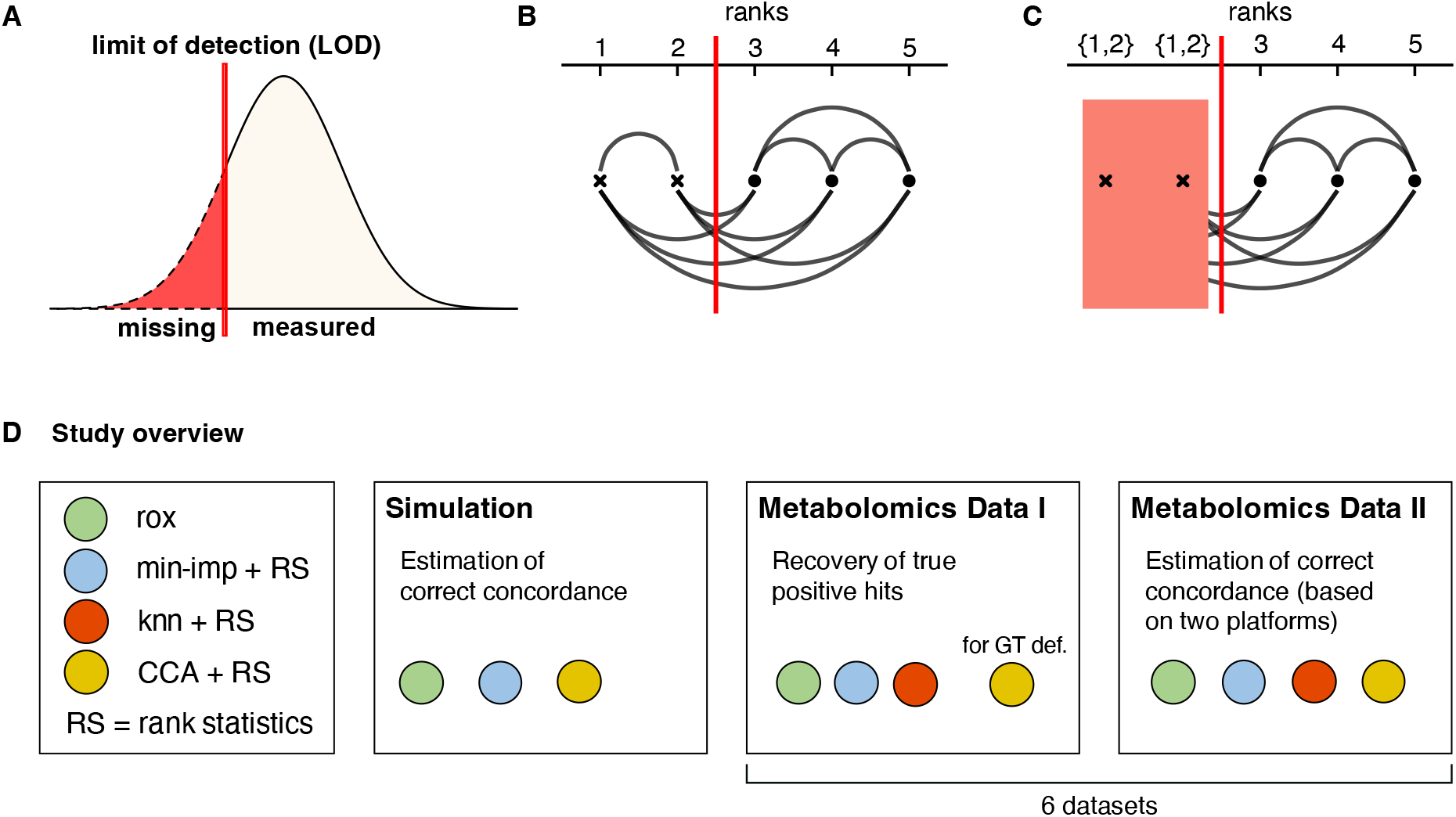
Limit of detection (LOD)-based missingness, statistical concept, and study overview. **A:** Schematic of a strict LOD effect on the distribution of a measurement. Values below the LOD (red line) will be reported as missing. **B:** Relative order of data points based on their true value. The red line indicates the theoretical LOD. **C:** Observed ordering of data points after LOD censoring. While observations below the LOD (red line) cannot be compared once they are censored, we still retain the information that all points below the LOD are lower than all points above the LOD. **D:** Overview of rox benchmarking. We assess the performance of our approach using an extensive simulation framework, followed by two test scenarios of ground truth recovery on a series of metabolomics datasets. “for GT def.” = used for ground truth definition.

A statistical method for the analysis of molecular data should take into consideration the above-mentioned issues and properties of missing values. First, it should make use of the fact that a missing value indicates a “low” abundance value, even if the precise numeric value is unknown. This allows to fully leverage the information available in the dataset. Ideally, the method should also work in the presence of a non-strict LOD mechanism. Second, in order to be applicable to a wide variety of molecular data, the method should be free of distributional assumptions and robust to outliers.

Existing statistical methods dealing with LOD-based missing values do not or only partially fulfill these requirements. The most popular approaches fall into one of three categories. (1) Missing values are simply deleted from the dataset, which is commonly referred to as complete case analysis (CCA) [6]. Since all samples with any missing values are removed, CCA often leads to a severe reduction of statistical power, especially when multivariate statistical methods are used. Moreover, if there is an enrichment of missing values in one of the analyzed groups, e.g., in sick individuals compared to healthy ones, CCA will substantially distort the statistical analysis and produce erroneous results [3]. (2) Imputation approaches where a full data matrix is reconstructed by replacing missing values with reasonable substitutes. “Minimum imputation” is a widely used approach that replaces missing values with the lowest observed value in the data, half of that value, or with a known LOD value [3]. Notably, this approach uses the information that missing values are low, but leads to a substantial distortion of the distribution of the analyte [7]. Other common approaches, such as k-nearest-neighbor (knn) imputation, use the correlation structure of the data to infer the original value [8]. These approaches do not use the LOD information and require a strong correlation structure among variables to work properly. (3) Statistical methods that directly incorporate the knowledge of the LOD effect, where missing values are treated as a “low” category. The approach published by [9] addresses the problem of LOD-based left-censoring in measurement data using methods from survival analysis, which we will prove later in this paper is equivalent to using rank statistics on minimum imputed data. Other approaches make specific assumption about the underlying data distribution (e.g., log-normal [10] or gamma [11]), and treat missing values as left-truncated data points from that respective distribution. While incorporating the LOD information, these methods also require strong assumptions on the overall data distribution, which might not be appropriate for certain data types.

Here we present *rox*, “rank order with missing values(X)”, a flexible, non-parametric approach for regression analysis of a dependent variable with missing values and continuous, ordinal, or binary explanatory variables. The core idea is to utilize the knowledge of missing values representing low concentrations due an LOD effect, without requiring any actual imputation steps. The approach is based on rank statistics related to Somer’s D and Kendall’s tau [12, 13, 14], and can be computed even with partially quantitative measurements (Figure 1B and C). Leveraging the properties of rank statistics, this framework is applicable to data from any distribution and is robust to outliers. Moreover, while the method relies on the assumption of an LOD effect in its core, it flexibly generalizes to data with other missingness mechanisms.

In this paper, we showcase the features of *rox* on simulated data and benchmark its performance on six real molecular datasets. We use metabolomics data, which is known to be heavily affected by LOD-based missingness and therefore constitutes an optimal test case for this approach. Notably, both for the simulated data and the real data, we define a ground truth for unbiased evaluation. Our analysis demonstrates the superiority of our approach over three of the most commonly used approaches in the field, namely complete case analysis (CCA), minimum imputation and knn-based imputation, coupled with rank-based statistical testing (Figure 1D). Our *rox* implementation is available as open source R package at https://github.com/krumsieklab/rox.

## 2 RESULTS

### 2.1 Simulation Results: Strict LOD

The *rox* method uses rank-based statistics to model measurements with limit of detection (LOD)-based missing value patterns, utilizing the information that absent data points represent low concentrations. It models the relationship between a measurement with missing values as the dependent variable and one or more continuous, ordinal, or binary explaining variables with no missing values. The approach furthermore implements a self-adjusting feature, which detects cases of non-LOD missingness, in which it switches to complete case analysis (CCA). A detailed mathematical derivation of the approach and its properties is provided in the METHODS section.

To show how *rox* performs at correctly recovering true concordance, we developed an extensive simulation framework with a known ground truth. The performance of *rox* was compared to that of complete case analysis (CCA) and regular concordance calculation after minimum imputation (min-imp). Note that k-nearest-neighbor (knn) imputation was omitted for this part, since it is only feasible in a multivariate setup, where simulation is dependent on various design choices and could easily be tweaked for a method to outperform the others. knn imputation will be evaluated based on real datasets later. In the first simulation, a single variable *Y* and a continuous outcome *X* were simulated with concordance *d* ranging between 0.55 and 0.85. These predefined concordance values represent the ground truth used to evaluate performance. Missing values were introduced into *Y* using a strict LOD mechanism, i.e., by setting all values below a given threshold to missing (Figure 2A). We simulated a variety of scenarios by ranging the proportion of missing values in *Y* from 0% to 90%. For each combination of true concordance and missing value proportion, we computed the concordance between *Y* and *X* by: (i) compute the *rox* statistic, (ii) imputing missing values in *Y* with minimum value imputation and performing regular concordance analysis, and (iii) only considering complete cases without missing values and performing regular concordance analysis. A large sample size of n = 10, 000 was chosen to ensure stable results. All simulations were repeated for smaller sample sizes, which yielded equivalent results (see Supplementary Figure 1).

**Figure 2:**
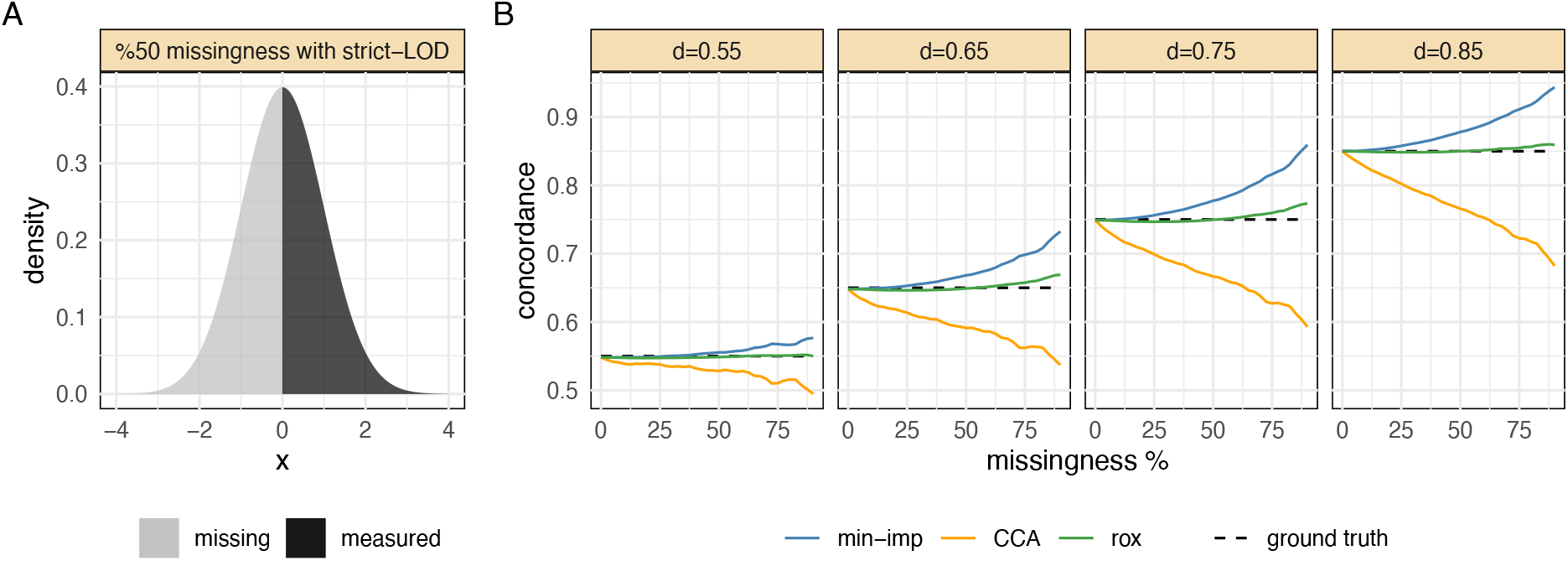
Simulation with strict LOD mechanism. **A**: Example distribution of a simulated variable with 50% missingness due to a strict LOD effect. **B**: *rox* outperformed CCA and minimum-imputation in recovering the true concordance across various ground truth values *d* and missingness fractions. Minimum imputation led to an overestimation of concordance between variable and outcome, while CCA resulted in underestimation.

The results of this first simulation demonstrated that *rox* was consistently better at retrieving the true concordance than its two competitors (Figure 2B). Minimum imputation generally led to an overestimation of concordance, while CCA led to an underestimation. With increasing proportions of missing values, these deviations increased substantially, while *rox* estimates remained stable and accurate. The same effect was observed across all values of true concordance values *d*.

### 2.2 Simulation Results: Probabilistic LOD

In the second simulation scenario, we evaluated the performance of *rox* in the case of a more realistic “probabilistic LOD” [3], where instead of a hard LOD threshold, the probability of a value being missing continuously increases with decreasing true abundance. A probabilistic LOD was simulated using a sigmoid probability density function that models the likelihood of a value being missing given its true value (Figure 3A). The shape of the sigmoid function was parametrized with a variable p*LOD*, which controls the type of missingness pattern in the data. *pLOD* =0 leads to missing at random (MAR) [15], while *pLOD* =1 generates a strict LOD effect. Therefore, higher values of *pLOD* lead to more prominent censoring effects (Figure 3B). In this scenario, we again simulated two random variables *Y* and *X* with true concordance values ranging from *d* = 0.55 to d = 0.85 and *pLOD* values ranging from 0 to 1. For this setup, the proportion of missing values in *Y* was fixed at 50%.

**Figure 3:**
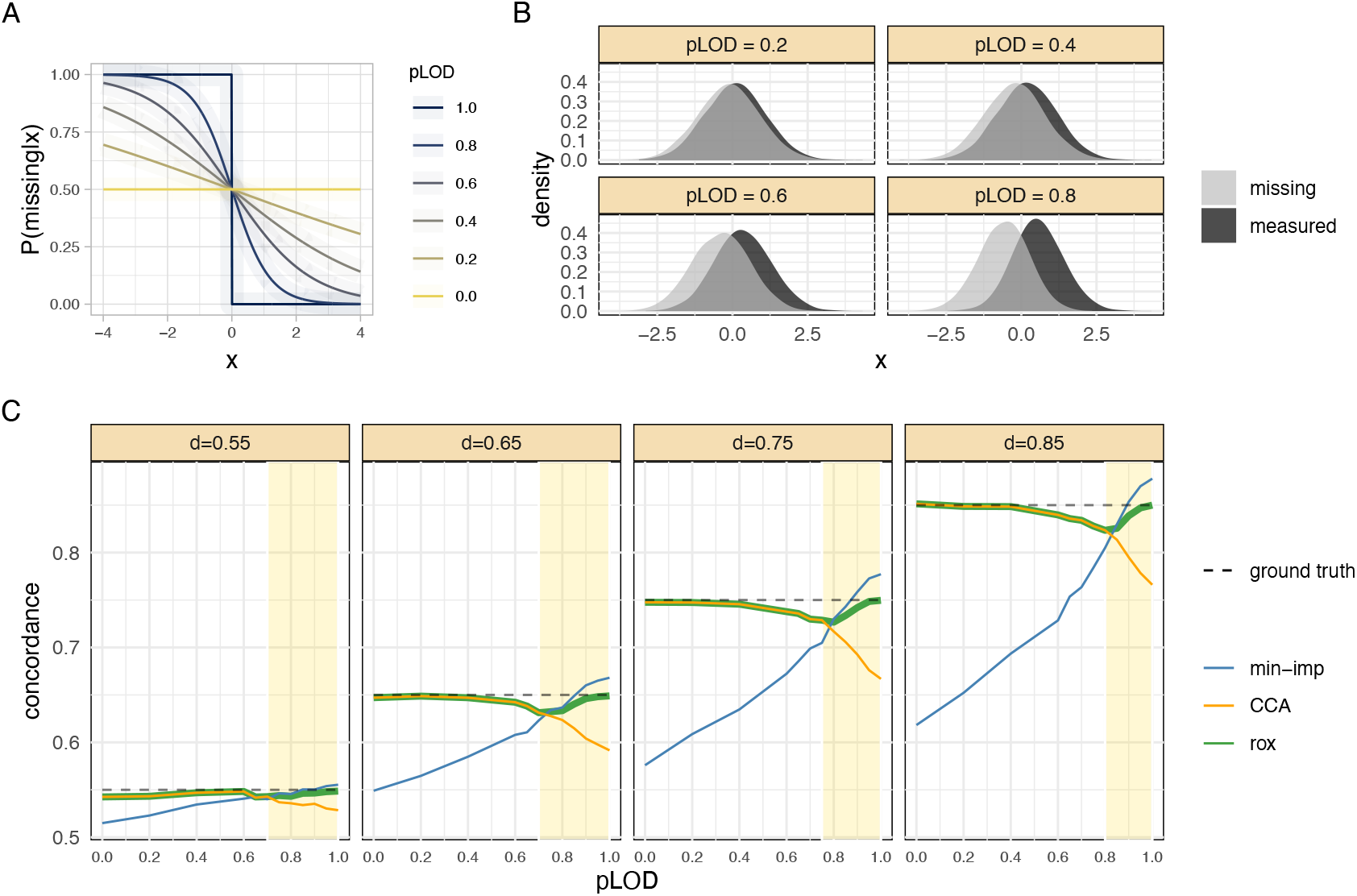
Simulation with probabilistic LOD mechanism. **A**: Probability function describing the likelihood of a value being missing as a function of its numerical value. A *pLOD* value of 1 results in a strict LOD effect, while pLod =0 results in values missing at random (MAR), and all values in between produce probabilistic LOD effects. **B**: Illustration of different missingness patterns induced by varying pLOD values. **C**: Data was simulated with different pLOD and concordance values, while the missingness percentage was kept at 50% in all scenarios. The yellow shaded area marks the region where our adaptive method automatically identified an active LOD effect and switched from CCA to *rox* analysis. Overall, *rox* outperformed minimum imputation and complete case analysis in all simulation settings.

The *rox* method again outperformed the competitor methods in all scenarios (Figure 3C) due to the adaptive nature of the model. For simulations with *pLOD* values below 0.7, *rox* determined that no sufficiently strong LOD effect was present and thus switched to complete case analysis (CCA). Minimum imputation, on the other hand, consistently underestimated concordance in the range of *d* between 0 and 0.7 due to its strict, implicit LOD assumption. For *pLOD* > 0.7, *rox* leveraged the left-censoring effect and consistently produced more accurate results than its competitors. In this high *pLOD* range, minimum imputation overestimated the concordance, and the performance of complete case analysis progressively deteriorated. This behavior was further exacerbated at increasing values of the true concordance *d*.

### 2.3 Simulation Results: Multivariable setting

In a third simulation, we investigated how the *rox* model performed in a multivariable setting for both strict-LOD and probabilistic-LOD scenarios. The multivariate setting is of particular interest when the inclusion of multiple variables in the same model is required, for example for covariate correction purposes. In this scenario, we simulated a continuous outcome, a covariate, and a variable of interest under various LOD and missingness settings and again compared *rox*’s performance with that of minimum imputation and knn imputation followed by regular concordance analysis (see Supplementary Figure 2). Results and conclusions were equivalent to the results of univariate simulations suggesting that *rox* recovered the ground truth better than competitor approaches.

### 2.4 Evaluation on Metabolomics Data: Recovering High-Confidence Hits

After evaluating the performance of *rox* in a simulation setting, we tested the approach in a real data scenario using published metabolomics datasets from a series of case-control studies (Table 1). In this case, we sought to determine how many true associations between metabolites and the respective study outcomes (e.g., disease status) *rox* could identify compared to its competitor methods.

**Table 1:** Overview of the metabolomics datasets.

| Cohort | Number of samples<br>(controls/cases) | Number of<br>metabolites | Phenotype | Specimen | Reference |
| --- | --- | --- | --- | --- | --- |
| QMDiab-Plasma (HD2) | 358<br>(177/181) | 758 | Type 2 Diabetes | Blood | [16] |
| QMDiab-Urine | 360<br>(174/186) | 891 | Type 2 Diabetes | Urine | [16] |
| QMDiab-Saliva | 330<br>(171/159) | 602 | Type 2 Diabetes | Saliva | [16] |
| BRCA | 132<br>(65/67) | 536 | Breast Cancer | Breast Tissue | [18] |
| RCC | 276<br>(138/138) | 877 | Kidney Cancer | Kidney Tissue | [19] |
| HAPO | 115<br>(67/48) | 49 | Hyperglycemia | Plasma | [17] |
| QMDiab-Plasma<br>Validation (HD4) | 292<br>(137/155) | 359 | Type 2 Diabetes | Plasma | [31, 32] |

Defining a ground truth in real datasets, however, is inherently difficult, since a list of true associations is usually not available. For our evaluation framework, we thus constructed a set of metabolite-outcome associations with high confidence of being actual true positives (HC-hits). This set was defined by combining the significant associations obtained from two statistical approaches that are well-suited to detect associations in data with antithetical missing values mechanisms. (1) CCA with Wilcoxon rank-sum test, which compares two sample groups by filtering out all samples with missing values. This test works well if values are missing at random and thus do not originate from an LOD effect but is generally underpowered since it entirely excludes missing values from the analysis. (2) Fisher’s exact test, which assess the proportion of missing values in one sample group versus the other, ignoring the actual numeric measurement values. In an LOD setting, this test works well for cases of extreme sample separation, for example when all samples in one of the comparison groups are low and fall below the LOD threshold. Notably, both approaches suffer from a substantial number of false negatives, since neither is ideally fit for the analysis of molecular data with missing values; however, both methods have very low false positive rates, meaning that the hits they identify are very likely to be correct.

In the following, we used the fraction of HC-hits that each method is able to retrieve as an evaluation metric. The analysis was performed on six datasets: Plasma, urine, saliva metabolomics from the QMDiab study [16], where the outcome was type-2-diabetes (T2D), a hyperglycemia study in pregnant women (HAPO) [17], with fasting plasma glucose (low FPG vs. high FPG) as outcome, and two tissue metabolomics datasets, one from breast tissue (BRCA) [18] and one from kidney tissue (RCC) [19], where the outcome was the origin of the sample (tumor or adjacent-normal tissue) of the sample. Across all datasets, *rox* outperformed or tied with the other methods in recovering HC-hits at various significance levels (Figure 4).

**Figure 4:**
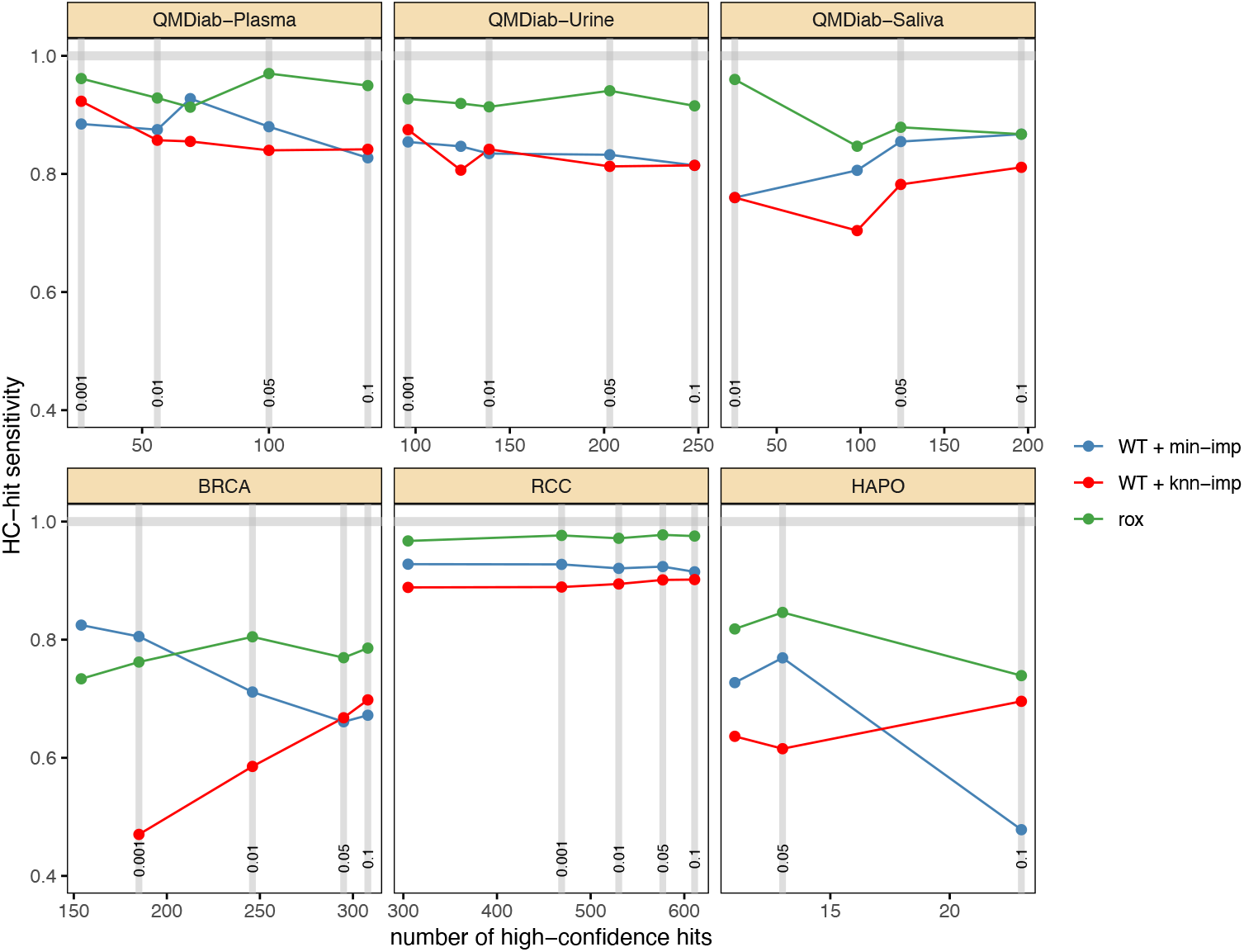
Recovery of high-confidence (HC) hits in six metabolomics datasets. The x-axis shows the number of HC-hits identified with the corresponding Bonferroni-adjusted p-value cut-off. The y-axis represents percentage of HC-hits that were identified, which is equivalent to a measure of sensitivity, at the respective cutoff. *rox* outperformed or tied with the two imputation approaches across all datasets and cutoffs.

### 2.5 Evaluation on Metabolomics Data: Multi-Platform Validation

A second line of validation on real data was performed on a dataset from the QMdiab study, where the same metabolites were measured in the same samples using two different metabolomics platforms. Specifically, we analyzed metabolites that were fully quantified (FQ) on one platform but were partially missing (PM) on the other platform. We used the concordance between the study outcome and the FQ metabolites without missing values as our ground truth and evaluated the performance of each method based on the consistency between this ground truth and the concordance estimate between the PM metabolite with missing values and the same outcome. In an ideal scenario, these two concordance values would be the same, indicating that the method recovered the correct value even in the presence of missing values. To analyze a sufficient number of FQ-PM metabolite pairs, we allowed up to 5% missing values in the FQ candidate and deleted those missing values in the subsequent analysis. If both platforms showed less than 5% missingness for a metabolite, we picked the one with the lower number of missing values as FQ metabolite, and the respective other measurement as PM metabolite.

Notably, missing values in these platforms are mostly due to a prominent LOD effect, which we confirmed by comparing missing and quantified values within the same metabolites across the two platforms (see Supplementary Figure 3). Thus, we expected this dataset to provide a favorable setting for minimum imputation, which assumes strict LOD. knn imputation, on the other hand, cannot impute values outside of the observed data distribution and is therefore unlikely to perform well in a strong LOD scenario [20].

Association analyses were performed between both FQ and PM metabolites and the respective QMdiab study outcomes (age, sex, BMI, and diabetes), using the *rox* test as well as regular association analysis with Wilcoxon rank-sum test after min-imp, knn-imp, and CCA. The PM-based concordance values were then compared with the ground truth concordance obtained from the corresponding FQ metabolite. An example of the results for age and PM metabolites with 20% or more missingness is shown in Figure 5A. This analysis was systematically repeated for varying fractions of missingness in the PM metabolite (see Figure 5B, and Supplementary Figure 4 for more detailed results). For all outcomes, *rox* was substantially more consistent than minimum-imputation, regardless of missingness percentage of the PM metabolite. Notably, the performance of minimum-imputation worsened with increasing missingness, while *rox*’s performance remained stable. *rox* outperformed knn-imputation specifically in the association with age, sex and BMI, while the two methods were mostly comparable for diabetes. Taken together, *rox* performed equivalently or better than knn-imputation across all scenarios. Similar comparisons were performed using multivariable *rox*, with analogous results (see Supplementary Figure 5).

**Figure 5:**
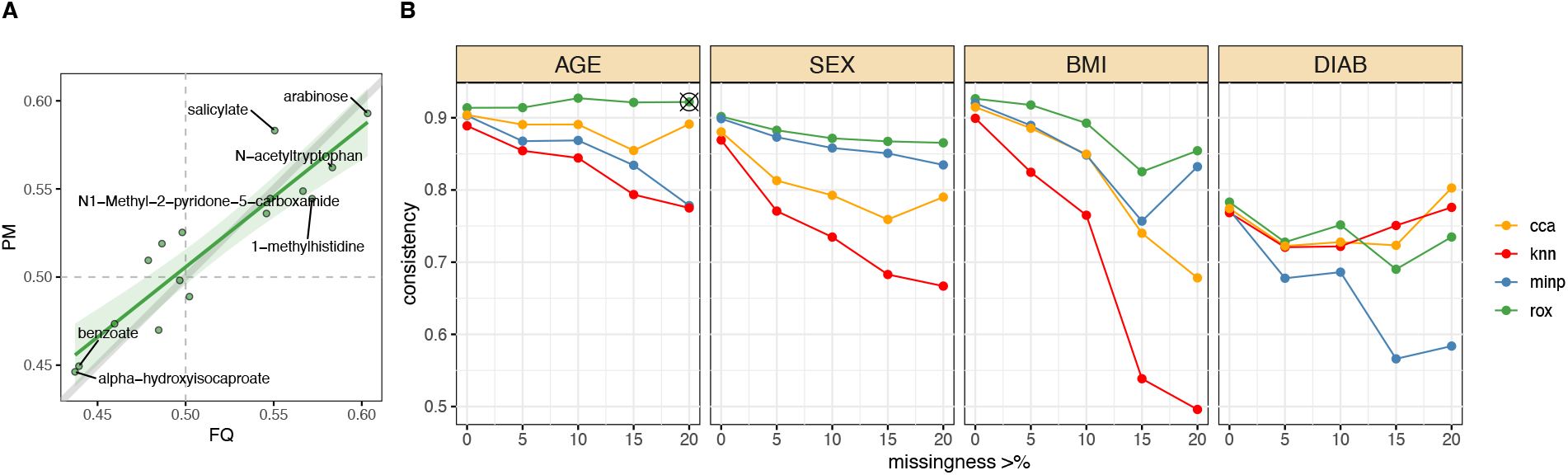
Validation on based two-platform comparison of fully quantified (FQ) and partially missing (PM) metabolites. Plasma metabolites from the same samples were measured on two different platforms. Consistency of concordance estimates between the FQ-based ground truth and the PM-based estimates was computed for all considered outcomes (age, BMI, diabetes (DIAB) and sex). **A:** Consistency across platforms for age associations calculated for PM metabolites with 20% or more missingness. Points distributed along the diagonal indicate consistent estimations between the two platforms. The gray line indicates the x=y axis, while the green line indicates a linear fit over the points of comparison. **B:** Systematic results for all four outcomes and varying levels of missingness in the PM metabolites. The x-axis indicates the minimum fraction of missing values per metabolite. The y-axis shows the Pearson correlation between the estimates across the two platforms. Results shown in panel A correspond to the point marked with the black cross. Across all missingness percentages, *rox* was more consistent compared to knn imputation and comparable to or better than minimum imputation.

## 3 DISCUSSION

This paper introduced *rox*, a novel statistical framework for datasets with missing values occurring due to a limit-of-detection (LOD) effect. In contrast to the more common approach of imputing missing values, which comes with various data analysis-related issues, *rox* directly utilizes the information that missing values have “low” concentrations. The non-parametric model is based on pairwise ranks and is thus robust to outliers. The method allows for multivariable modeling and can be used for any quantitative or semiquantitative measurements. Importantly, while *rox* is inherently designed for data with an LOD effect, it also works with less strict, blurry LOD-based data, or even when the values are missing at random, in which case it automatically switches to complete case analysis. Using a simulation framework as well as metabolomics datasets from various sample types with different outcomes, we systematically demonstrated the superiority of our method over other approaches that are commonly used in the field. Specifically, *rox* showed higher accuracy in reconstructing the underlying, true concordance values, and displayed higher statistical power retrieving associations with study outcomes. Notably, while most other studies on real data artificially introduce missing values to evaluate the performance of their statistical approach (e.g., [8, 21]), we here relied on two data-driven frameworks to define a ground truth for a more realistic evaluation.

In conclusion, we recommend using *rox* for any dataset where an LOD effect can be suspected, even if the effect is not strict. The LOD assumption commonly applies to metabolomics data, as shown in this paper, but has also been described in data with similar dropout mechanisms, such as proteomics data [22], glycomics data [23], and microbiomics data [24].

## 4 METHODS

### 4.1 *rox* core model

*rox* is inspired by the ranking-based, non-parametric correlation measure *concordance index* or c-index [25], which is equivalent to an ROC-AUC with a binary outcome [26]. Let S = {(*x*_*1*_, *y*_*1*_), …, (*x*_*n*_, *y*_*n*_)} be a set of n observations of two random variables *X* and *Y*. A pair of observations ⟨ *i, j* ⟩= {(*x*_*i*_, *y*_*i*_), (*x*_*j*_, *y*_*j*_)} is said to be *concordant* if the pairwise ranking of (*x*_*i*_, *x*_*j*_) and (*y*_*i*_, *y*_*j*_) is the same, i.e., if sgn (*x*_*i*_− *x*_*j*_) = sgn (*y*_*i*_− *y*_*j*_); otherwise it is said to be *discordant*. The c-index is then defined as the number of concordant pairs over the number of all pairs 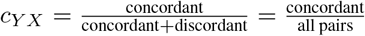 [25]. Note that in *c*_*YX*_, ties in *Y* are dropped from the calculation, whereas in *X* are counted as 0.5, i.e., neither concordant nor discordant.

The *rox* statistic is an extension of this c-index concept to left-censored data. This type of data occurs, for example, when all values below a certain threshold (the *Limit Of Detection*, or LOD) are returned as missing. Based on this LOD assumption, we know that any missing value is lower than any measured value in the data. Importantly, a missing and a measured value can thus still be ranked and are hence *comparable*, while two missing values have no known order and are *non-comparable*. The *rox* method assesses the fraction of concordant pairs only in relation to the *comparable pairs*, 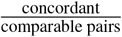, a concept which is also used in survival analysis [25]. In the following, we formulate a concordance-based test on comparable pairs.

Let *Y* now be a left-censored random variable with LOD-based missing values and let *X* be an outcome of interest to be associated with *Y*. As outlined above, any pair of observations ⟨ *i, j* ⟩ where at least one of *y*_*i*_ or *y*_*j*_ is non-missing constitutes a comparable pair, since we know that the LOD requires, by design, any non-missing value to be larger than all missing values (see also, Figure 1B and C). Let *π* = {⟨*i, j*⟩ | *y*_*i*_ ≠ NA or *y*_*j*_ ≠NA} be the set of these comparable pairs, where NA represents a missing value, and let (*π*) be the number of concordant pairs in *π* given *X* and *Y*. The non-parametric *rox* coefficient of concordance between variables *X* and *Y*, with *Y* subject to LOD-based missingness, can be formulated as follows:

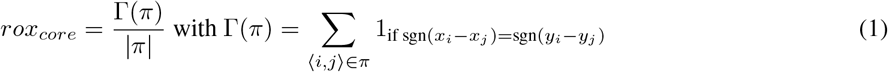

where |*π* | is the set size of *π*, i.e., the total number of comparable pairs.

In general, the sgn operator is not defined for missing values. However, under strict-LOD assumptions, any missing value in *Y* will be lower than all non-missing values. This means that sgn (*y*_*i*_ − *y*_*j*_) is always defined in this framework, even when either *y*_*i*_ or *y*_*j*_ is missing. Note that, similar to the c-index, the *rox* statistic represents the probability of concordance between *X* and *Y*, and it can take values in the interval [0,1], where 0.5 indicates random ordering, 1 represents perfect concordance, and 0 represents perfect discordance.

### 4.2 Debiased, weighted *rox* model

In Eq.(1), pairs where both *y*_*i*_ and *y*_*j*_ are missing constitute non-comparable pairs, and are excluded from the concordance estimation. However, ignoring these pairs leads to an overall overestimation of positive concordance (> 0.5) and an underestimation of negative concordance (< 0.5); see remark at the bottom of Supplementary Text 2. Note that the 0.5 cut point is due to the scale of concordance between 0 and 1, where values above 0.5 indicate positive correlation and values below 0.5 represent negative correlation.

To address this problem, we propose a strategy to de-bias the *rox* coefficient by down-weighting the contribution of missing observations to the overall concordance. To this end, we split all comparable pairs from Eq.(1) into two distinct sets: *π*_*b*_ (bridge pairs), which includes pairs of observations where either *y*_*i*_ or *y*_*j*_ is missing, and *π*_*1*_, which includes pairs where both *y*_*i*_ and *y*_*j*_ are non-missing (see Supplementary Figure 6). This way, all comparable pairs are partitioned as *π* = *π*_*b*_ ⋃*π*_*1*_.

With this formulation, we can now introduce a weight parameter *p* to control the contribution of the pairs with missing values, *π*_*b*_, to the overall *rox* statistics as:

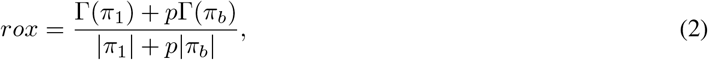

where 0 ≤ *p* ≤ 1 (see Supplementary Text 1 for a detailed derivation). Setting *p* =1 leads to the original formulation from Eq.(1), which is based on a strict LOD assumption, whereas *p* =0 reduces the test statistics to a non-parametric complete case analysis, ignoring the contribution of all pairs including any missing values.

In general, if *n*_*0*_ is the number of missing values, n_*1*_ is the number of non-missing values, and *n* = *n*_*0*_ + *n*_*1*_ is the total number of observations, a higher fraction of missing values *n*_*0*_/*n* will introduce more bias in the concordance estimation and will hence require a lower value of the weight p to de-bias the estimate. In Supplementary Text 3, we demonstrate that the concordance can be effectively debiased by using the weight factor *p* = *n*_*1*_/*n*. With this new expression for p, Eq.(2) becomes:

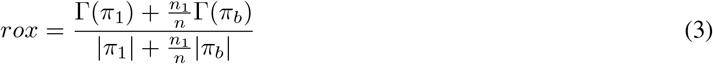

### 4.3 Self-adjusting *rox* for partial LOD and non-LOD

The weighted formulation in Eq.(3) assumes that missingness in *Y* occurs due to a strict LOD threshold, i.e., that *all* values below the LOD threshold will be missing and *all* values above the threshold will be present. However, in many real data scenarios, missingness patterns occur on a continuum [3], from a strict LOD mechanism, to a more probabilistic setting, where lower values have a higher likelihood of being missing, all the way to missing-at-random (MAR). For the *rox* statistic, non LOD-based missingness constitutes a source of bias that will affect the estimation of the true concordance. For cases where the missingness pattern is only marginally due to LOD or even LOD-independent, ignoring the missing values and switching to a complete case analysis is more appropriate.

We thus formulated a self-adjusting version of *rox*. First, we estimate whether the missingness pattern in the data is consistent with an LOD assumption. Let d_*1*_ = (*π*_*1*_)/ |*π*_*1*_| and d_*b*_ = (*π*_*b*_)/ |*π*_*b*_ |be the concordances of pairs with no missing values and pairs with one missing value, respectively. Under strict LOD, which corresponds to a left-truncation of the data distribution, if the true concordance is larger than 0.5, then it holds that *d*_*1*_ < *d*_*b*_, while for concordance values less than 0.5 it holds that *d*_*b*_ < *d*_*1*_ (see Supplementary Text 2 for proof). For simplicity, we only describe the positive concordance case here; the negative concordance case can be derived analogously.

For any random variable *Y*, we can assess whether the LOD assumption is violated by checking whether *d*_*1*_ < d_*b*_. If the inequality holds, *rox* concordance is estimated using Eq.(3); if it does not, *p* in Eq.(2) is set to zero, removing all missing observations from the analysis and effectively computing concordance based only on observations with no missing values, reducing the approach to a complete-case-analysis (CCA).

Thus, the final formulation of *rox* for positive concordance is:

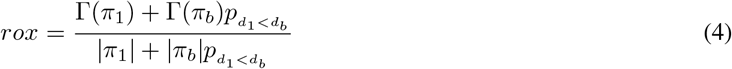

where 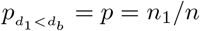 as in Eq.(3) if *d*_*1*_ < *d*_*b*_, and 0 otherwise.

### 4.4 *Rox*-based semi-parametric multivariable model

The *rox* model handles one-to-one relations between two variables. In this section, we extend the approach to a semi-parametric, multivariable modeling framework which allows to model relations between one variable with missing values and multiple variables.

The proposed extension is obtained via multivariable modeling of the concordance probabilities with an exponential link function [27]. Let Y be a metabolite measurement with missing values and **X** = {*X*_*1*_, *X*_*2*_, …, *X*_*k*_} be k different variables of interest. First, we define the likelihood of concordance for a single pair of observations ⟨ *i, j*⟩ as 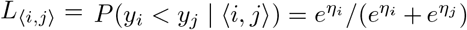 where *η*_*i*_ = *β*_*1*_*x*_*i1*_ + *β*_*2*_*x*_*i2*_ + … + *β*_*k*_*x*_*ik*_ is the standard linear predictor function for sample *i* and *β* = { *β*_*1*_, *β*_*2*_, …, *β*_*k*_} is the vector of the corresponding regression coefficients. The log-likelihood 𝓁 (*π*) of the associated joint probability of all realized pairwise rankings in *π* can then be formulated as the product of all individual likelihoods:

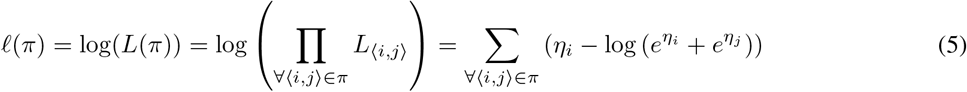

Similar to the univariate case, Eq.(5) ignores pairs of observations where both *Y* values are missing. However, ignoring these observations leads to a biased estimate of the concordance probability. Also in this case, we can therefore debias the model by down-weighing the contribution of missing values and accounting for non-LOD scenarios. We again partition the observation pairs in *π* into those between two observations with no missing value *π*_*1*_ and those where one of the two observations is missing *π*_*b*_ (i.e. *π* = *π*_*1*_ ⋃ *π*_*b*_), introduce a weight p to control the contribution of pairs with missing values, and check for the violation of LOD assumption based on the *d*_*1*_ < *d*_*b*_ inequality:

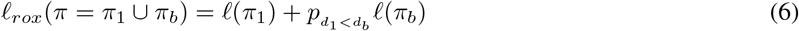

where *p* = *n*_*1*_/*n* as derived in Supplementary Text 3. All *β* coefficients are fitted using a maximum likelihood estimation (MLE) approach, based on a FORTRAN implementation for concordance regression we adapted from [27]. The overall concordance of the model is then calculated by computing the *rox* statistic from Eq.(4) between *Y* and the fitted score 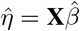.

### 4.5 Hypothesis testing

In the following, we derive a hypothesis test for the univariate version of *rox*, assessing whether *H*_*0*_ : *rox*(*Y, X*)= 0.5 can be rejected. Under the null hypothesis of independence of *Y* and *X*, the distribution of the quantity 2 × *rox* − 1 has an expected value of zero. A significance test for *rox* can be obtained via: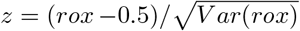, and the corresponding p-value can be calculated via a z-test. The variance of the concordance can be estimated in two different ways: (1) Using an equivalent time-dependent Cox model (*cvar*) [26] or (2) through an unbiased infinitesimal jackknife variance estimator (*ivar*) [28]. As pointed out by Therneau et al. [28], *cvar* is an unbiased estimator for *d* ≠ 0.5, while it overestimates the variance if *d* ≠ 0.5. On the other hand, *ivar* is unbiased for *d* ≠ 0.5, but underestimates the variance if *d* is close to 0.5. Taking these findings into consideration, we calculate the p-value of the estimated *rox* statistics based on the average of these two variance estimates, namely 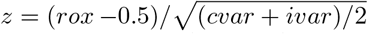. This approach was inspired by [29], where overestimated and underestimated variances are averaged to get a better estimate.

In a multivariable setting, hypothesis testing for the overall model is performed as described in the previous paragraph with *H*_*0*_ : *rox*(*Y*, **X** *β*)= 0.5, where {*β* = *β*_*1*_, *β*_*2*_, …, *β*_*k*_ } are the regression coefficients, and **X***β* is the linear predictor of the model. Furthermore, to test significance of individual variables in the model, we can use the coefficients in the proposed semi-parametric model. In this case, the null hypothesis is defined based on the coefficients: *H*_*0*_ : *β*_*i*_ =0 for variable *i*. To assess significance, we used the implementation of [27] to estimate coefficients and standard errors to calculate a Wald’s test [30].

### 4.6 Simulation framework

Two continuous variables *Y* and *X* with a predefined concordance values were simulated. However, concordance cannot be directly parameterized and needs to be determined empirically. Here, we generated the desired concordance by tuning an association parameter as follows: The variable *X* and a noise term ϵ were first sampled from a standard normal distribution. *Y* was then defined as *Y* = X + ϵ, where determines the association between *Y* and *X*. Larger values of lead to lower concordance between the two variables. We ranged from 0 to 0.7 in steps of 0.01 until the desired concordance *d*(*Y, X*) was reached.

For the large sample size simulation, we generated a total of *n* = 10, 000 samples. For the small sample size simulation, we first generated a large dataset of *n* = 1, 000, 000 samples, from which 100 random samples were drawn 1, 000 times.

In the multivariable case, we simulated a variable *Y*, an outcome of interest *X* and a covariate *Z*. Correlations between *X, Y, Z* were simulated as follows: *X* was sampled from a normal distribution. A correlated covariate *Z* is simulated as Z = X + *ϵ*_*z*_, with *ϵ*_*z*_ being a normal distribution. *Y* was sampled as *Y* = *X* + *Z* + 3ϵ_*y*_, with ϵ_*y*_ again being a normally distributed error term.

### 4.7 Metabolomics datasets

To illustrate the performance of *rox* on real data, we analyzed a total of seven previously published metabolomics datasets (Table 1). For the QMDiab plasma validation cohort (HD4), only samples and metabolites overlapping with the HD2 platform were considered. Except for the HAPO dataset, for which only preprocessed data were available, all datasets were preprocessed using the R package maplet [33] as follows: Prior to statistical analysis, raw peak intensities were normalized using the probabilistic quotient approach [34], using only metabolites with less than 20% missing values to generate the reference sample. Normalized metabolite values were subsequently *log*_*2*_ transformed. The following imputation step was applied for all datasets. For each cohort, two additional data matrices were generated: one where missing values were imputed using the minimum value per metabolite, and one where missing values were imputed using knn-based imputation with 10 neighbors and variable pre-selection based on pairwise correlation (threshold of 0.2) according to [3].

## References

[1] Liang Jin et al. “A comparative study of evaluating missing value imputation methods in label-free proteomics”. In: Scientific reports 11.1 (2021), pp. 1–11.

[2] Huang Lin and Shyamal Das Peddada. “Analysis of microbial compositions: a review of normalization and differential abundance analysis”. In: NPJ biofilms and microbiomes 6.1 (2020), pp. 1–13.

[3] Kieu Trinh Do et al. “Characterization of missing values in untargeted MS-based metabolomics data and evaluation of missing data handling strategies”. en. In: Metabolomics 14.10 (Sept. 2018), p. 128. ISSN: 1573-3890. DOI: 10.1007/s11306-018-1420-2. URL: https://doi.org/10.1007/s11306-018-1420-2(visited on 11/09/2020).

[4] Karsten Suhre et al. “Human metabolic individuality in biomedical and pharmaceutical research”. In: Nature 477.7362 (2011), pp. 54–60.

[5] Gregory B. Gloor et al. “Microbiome Datasets Are Compositional: And This Is Not Optional”. In: Frontiers in Microbiology 8 (Nov. 2017). DOI: 10.3389/fmicb.2017.02224. URL: https://doi.org/10.3389/fmicb.2017.02224.

[6] Ian R. White and John B. Carlin. “Bias and efficiency of multiple imputation compared with complete-case analysis for missing covariate values”. In: Statistics in Medicine 29.28 (Sept. 2010), pp. 2920–2931. DOI: 10.1002/sim.3944. URL: https://doi.org/10.1002/sim.3944.

[7] Dennis R. Helsel. “Fabricating data: How substituting values for nondetects can ruin results, and what can be done about it”. In: Chemosphere 65.11 (Dec. 2006), pp. 2434–2439. DOI: 10.1016/j.chemosphere.2006.04.051. URL: https://doi.org/10.1016/j.chemosphere.2006.04.051.

[8] O. Troyanskaya et al. “Missing value estimation methods for DNA microarrays”. In: Bioinformatics 17.6 (June 2001), pp. 520–525. DOI: 10.1093/bioinformatics/17.6.520. URL: https://doi.org/10.1093/bioinformatics/17.6.520.

[9] Dennis R Helsel et al. Nondetects and data analysis. Statistics for censored environmental data. Wiley-Interscience, 2005.

[10] Lawrence H. Moulton and Neal A. Halsey. “A Mixture Model with Detection Limits for Regression Analyses of Antibody Response to Vaccine”. In: Biometrics 51.4 (Dec. 1995), p. 1570. DOI: 10.2307/2533289. URL: https://doi.org/10.2307/2533289.

[11] D. B. Richardson. “Effects of Exposure Measurement Error When an Exposure Variable Is Constrained by a Lower Limit”. In: American Journal of Epidemiology 157.4 (Feb. 2003), pp. 355–363. DOI: 10.1093/aje/kwf217. URL: https://doi.org/10.1093/aje/kwf217.

[12] M. G. Kendall. “Rank and Product-Moment Correlation”. In: Biometrika 36.1/2 (1949). Publisher: [Oxford University Press, Biometrika Trust], pp. 177–193. ISSN: 0006-3444. DOI: 10.2307/2332540. URL: https://www.jstor.org/stable/2332540(visited on 11/08/2020).

[13] Roger Newson. “Parameters behind “nonparametric” statistics: Kendall’s tau, Somers’ D and median differences”. In: The Stata Journal 2.1 (2002), pp. 45–64.

[14] Robert H Somers. “A new asymmetric measure of association for ordinal variables”. In: American sociological review (1962), pp. 799–811.

[15] Donald B. Rubin. “Inference and missing data”. en. In: Biometrika 63.3 (Dec. 1976). Publisher: Oxford Academic, pp. 581–592. ISSN: 0006-3444. DOI: 10.1093/biomet/63.3.581biomet/article/63/3/581/270932. URL: https://academic.oup.com/ (visited on 11/09/2020).

[16] Kieu Trinh Do et al. “Phenotype-driven identification of modules in a hierarchical map of multifluid metabolic correlations”. en. In: npj Systems Biology and Applications 3.1 (Sept. 2017). Number: 1 Publisher: Nature Publishing Group, pp. 1–12. ISSN: 2056-7189. DOI: 10.1038/s41540-017-0029-9. URL: https://www.nature.com/articles/s41540-017-0029-9(visited on 11/09/2020).

[17] Denise M Scholtens et al. “Metabolomics reveals broad-scale metabolic perturbations in hyperglycemic mothers during pregnancy”. In: Diabetes care 37.1 (2014), pp. 158–166.

[18] Atsushi Terunuma et al. “MYC-driven accumulation of 2-hydroxyglutarate is associated with breast cancer prognosis”. In: The Journal of clinical investigation 124.1 (2014), pp. 398–412.

[19] A Ari Hakimi et al. “An integrated metabolic atlas of clear cell renal cell carcinoma”. In: Cancer cell 29.1 (2016), pp. 104–116.

[20] Lorenzo Beretta and Alessandro Santaniello. “Nearest neighbor imputation algorithms: a critical evaluation”. In: BMC medical informatics and decision making 16.3 (2016), pp. 197–208.

[21] Daniel J Stekhoven and Peter Bühlmann. “MissForest—non-parametric missing value imputation for mixed-type data”. In: Bioinformatics 28.1 (2012), pp. 112–118.

[22] Yuliya Karpievitch et al. “A statistical framework for protein quantitation in bottom-up MS-based proteomics”. In: Bioinformatics 25.16 (2009), pp. 2028–2034.

[23] Gerald W Hart and Ronald J Copeland. “Glycomics hits the big time”. In: Cell 143.5 (2010), pp. 672–676.

[24] Justin D Silverman et al. “Naught all zeros in sequence count data are the same”. In: Computational and structural biotechnology journal 18 (2020), pp. 2789–2798.

[25] F. E. Harrell et al. “Evaluating the yield of medical tests”. eng. In: JAMA 247.18 (May 1982), pp. 2543–2546. ISSN: 0098-7484.

[26] Terry Therneau and Elizabeth Atkinson. Concordance. en. vignette of survival package. Sept. 2020. URL: https://cran.r-project.org/web/packages/survival/vignettes/concordance.pdf.

[27] Daniela Dunkler, Michael Schemper, and Georg Heinze. “Gene selection in microarray survival studies under possibly non-proportional hazards”. en. In: Bioinformatics 26.6 (Mar. 2010). Publisher: Oxford Academic, pp. 784–790. ISSN: 1367-4803. DOI: 10.1093/bioinformatics/btq035com/bioinformatics/article/26/6/784/244757. URL: https://academic.oup.(visited on 11/09/2020).

[28] Terry M Therneau and David A Watson. “The concordance statistic and the Cox model”. en. In: Department of Health Science Research Mayo Clinic Technical Report.85 (2017), p. 18.

[29] Stefan Wager, Trevor Hastie, and Bradley Efron. “Confidence intervals for random forests: The jackknife and the infinitesimal jackknife”. In: The Journal of Machine Learning Research 15.1 (2014), pp. 1625–1651.

[30] Abraham Wald. “Tests of statistical hypotheses concerning several parameters when the number of observations is large”. In: Transactions of the American Mathematical society 54.3 (1943), pp. 426–482.

[31] Kieu Trinh Do et al. “MoDentify: phenotype-driven module identification in metabolomics networks at different resolutions”. In: Bioinformatics 35.3 (2019), pp. 532–534.

[32] Dennis O Mook-Kanamori et al. “1, 5-Anhydroglucitol in saliva is a noninvasive marker of short-term glycemic control”. In: The Journal of Clinical Endocrinology & Metabolism 99.3 (2014), E479–E483.

[33] Kelsey Chetnik et al. “maplet: An extensible R toolbox for modular and reproducible metabolomics pipelines”. In: Bioinformatics 38.4 (2022), pp. 1168–1170.

[34] Frank Dieterle et al. “Probabilistic quotient normalization as robust method to account for dilution of complex biological mixtures. Application in 1H NMR metabonomics”. In: Analytical chemistry 78.13 (2006), pp. 4281– 4290.

[35] William H Kruskal. “Ordinal measures of association”. In: Journal of the American Statistical Association 53.284 (1958), pp. 814–861.

[36] Kendall, M.G. Rank Correlation Methods. London: Charles Griffin and Co, 1948.

[37] Gregory A Fredricks and Roger B Nelsen. “On the relationship between Spearman’s rho and Kendall’s tau for pairs of continuous random variables”. In: Journal of statistical planning and inference 137.7 (2007), pp. 2143– 2150.

