## Supplementary Methods and Figures for "*rox*: A statistical model for regression with missing values"

### Supplementary Materials

##### Supplementary Text 1: Derivation of weighted formulation

Starting from the core $rox$ formulation in Eq.(1)

$$rox_{core} = \frac{\Gamma(\pi)}{|\pi|} \text{ where } \Gamma(\pi) = \sum_{\langle i,j \rangle \in \pi} 1_{\text{if } \text{sgn}(x_i - x_j) = \text{sgn}(y_i - y_j)} \quad (1)$$

Let $w_{ij}$ be the weight of pair $\langle i, j \rangle$, and $\Gamma(\pi, w)$ be the sum of the weights of concordant pairs in $\pi$. We first introduce a variation of the $rox$ statistic that includes a contribution weight for each pair of observations as:

$$rox(w) = \frac{\sum_{\langle i,j \rangle \in \pi} w_{ij} 1_{\text{if } \text{sgn}(x_i - x_j) = \text{sgn}(y_i - y_j)}}{\sum_{\langle i,j \rangle \in \pi} w_{ij}} = \frac{\Gamma(\pi, w)}{\sum_{\langle i,j \rangle \in \pi} w_{ij}}, \quad (1.1)$$

Notably, if all weights are unitary, i.e., $w_{ij} = 1 \forall i, j$, this expression simplifies to the core formulation in Eq.(1).

We now want to derive a formulation that allows to control the overall contribution of missing values to the statistic. Under an LOD assumption, we can divide all comparable pairs $\pi$ in two sets of pairs: one set of pairs which include a missing value, referred to as $\pi_b$, and one set of pairs consisting of only complete cases, referred to as $\pi_1$. This way, $\pi = \pi_1 \cup \pi_b$, and $w = w_1 \cup w_b$, from which follows:

$$rox(w) = \frac{\Gamma(\pi, w)}{\sum_{\langle i,j \rangle \in \pi} w_{ij}} = \frac{\Gamma(\pi_1, w_1) + \Gamma(\pi_b, w_b)}{\sum_{\langle i,j \rangle \in \pi_1} w_{ij} + \sum_{\langle i,j \rangle \in \pi_b} w_{ij}} \quad (1.2)$$

We assign a weight equal 1 to all the pairs with complete observations, and set the same weight $p$ with $0 \leq p \leq 1$ to all pairs that contain a missing value. That is, $w_{ij} = 1 \; \forall i, \forall j$, and $w_{ij} = p$ otherwise. This results in Eq.(2):

$$rox(p) = \frac{\Gamma(\pi_1) + p\Gamma(\pi_b)}{|\pi_1| + p|\pi_b|} \quad (2)$$

##### Supplementary Text 2: Proof - LOD leads to $d_b > d > d_1$

Under the LOD assumption, $rox$ accepts missing values as left-censored values and offers an unbiased estimate of concordance. The framework has a self-adjustment feature, which checks whether there is evidence that measurements violate this LOD assumption. In that case, $rox$ switches off the left-censored formulation, since non-LOD data would only further bias the estimate, see Eq.(3), Eq.(4) in the main manuscript. Whether the data is under an LOD effect is assessed by evaluating whether the inequality $d_b > d > d_1$ holds (as defined in main manuscript). If the inequality does not hold, $rox$ concludes a violation of the LOD assumption, and switches to a complete case analysis (CCA), ignoring all missing values. In this supplement, under some mild assumptions, we will prove that if a strict-LOD exists, then $d_b > d > d_1$ must hold. We will first introduce and quantities and notations that will be useful later on, and will then show the proof for the inequality.

##### Variables and Nomenclature

Let $X$ and $Y$ be two random variables from a bivariate distribution $(X, Y)$, and let $d = d_{YX}$ be the overall concordance between $Y$ and $X$. Let $S$ be the set of all observations, and let us partition $S$ into two disjoint subsets $s = \{s_1 \subsetneq S, s_2 \subsetneq S \mid S = s_1 \cup s_2,\ s_1 \cap s_2 = \emptyset\}$. Based on this definition, we can partition all pairwise rankings $\pi$ into three distinct sets of pairs $\pi_1$, $\pi_2$, and $\pi_b$, where $\pi_1$ includes all the pairs within $s_1$, $\pi_2$ includes all the pairs within $s_2$, and $\pi_b$ includes the pairs between the two sets, i.e., the "bridge pairs" (Supplementary Figure 7 below). Note that $\pi = \pi_1 \cup \pi_b \cup \pi_2$. Furthermore, let $(X^{(s_i)}, Y^{(s_i)})$ denote the observations of $X$ and $Y$ relative to a specific set $s_i \in \{s_1, s_2\}$. In this setting, $d_1 = d_{Y^{(s_1)}X^{(s_1)}}$ is the concordance between $X$ and $Y$ within $s_1$, and hence over the pairs $\pi_1$. Likewise $d_2 = d_{Y^{(s_2)}X^{(s_2)}}$ is the concordance between $X$ and $Y$ within $s_2$, over the pairs in $\pi_2$. Finally, we can define $d_b$ as the concordance between $X$ and $Y$ over the set of bridge pairs $\pi_b$.

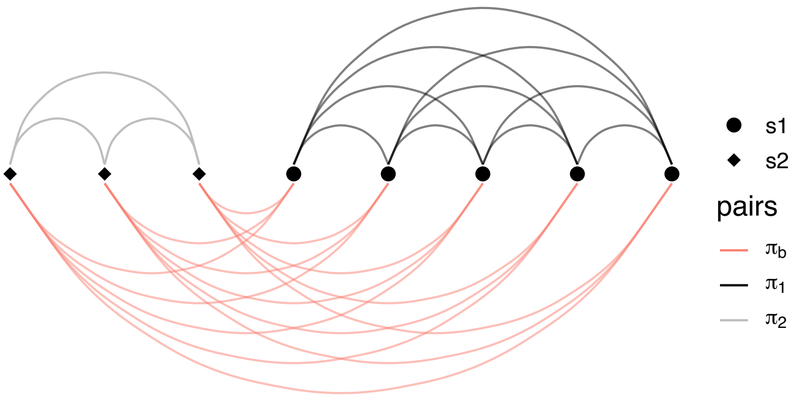

Supplementary Figure 7: By separating the set of all observations $S$ into two non overlapping subsets $s_1$ and $s_2$, we define three distinct sets of pairs $\pi_1$, $\pi_2$ and $\pi_b$, where $\pi_1$ and $\pi_2$ include pairs within $s_1$ and within $s_2$, respectively, and $\pi_b$ includes pairs constituted by one observation from $s_1$ and one observation from $s_2$.

In all following derivations, we will assume the true concordance between $X$ and $Y$ to be positive, i.e., $d_{YX} > 0.5$. For negative concordance, results can be derived analogously by noting that $d_{YX} < 0.5$ corresponds to $d_{Y(-X)} > 0.5$. Therefore, simply substituting $X$ with $-X$ in all following concordance derivations is sufficient to obtain the results for the negative concordance case.

##### Lemmas

**Lemma 1:** Greiner's relation [35] states that for a bivariate normal distribution of random variables $X$ and $Y$, Kendall's $\tau$ and Pearson correlation $\rho$ are related by $\rho_{XY} = sin(\frac{\pi}{2}\tau_{XY})$. Thus, solving this for $\tau$ yields

$$\tau_{XY} = G(\rho_{XY}) \text{ where } G(x) = \frac{2}{\pi}arcsin(x) \tag{7}$$

**Lemma 2:** Given two bivariate normal random variables $X$ and $Y$ and a subset of observations $s \subsetneq S$ such that $Var[X(S)] > Var[X(s)]$, the Kendall's correlation between $X$ and $Y$ based on the subset of observations $s$ is smaller than the correlation based on the full set of observations $S$.

Given a bivariate normal distribution of random variables $X$ and $Y$, suppose that $Y = \beta X + \epsilon$, where $\beta$ is the linear regression coefficient and $\epsilon$ is a Gaussian error term. The variance of $Y$ can be written in terms of the variance of $X$ as $Var[Y] = \beta^2 Var[X] + Var[\epsilon]$. In this setting, Pearson correlation $\rho$ is related to the linear model's explained variance $R^2$ by $\rho^2 = R^2$, which can be written as

$$\rho^2 = R^2 = \frac{\beta^2 Var[X]}{\beta^2 Var[X] + Var[\epsilon]} \tag{8}$$

The right-hand side of Eq.(8) goes to zero as $Var[X] \to 0$. This means that the magnitude of Pearson correlation $|\rho|$ decreases as $Var[X] \to 0$. Now let $s$ be a subset from the population of observations $S$, $s \subsetneq S$, such that $XY^{(s)} = (X^{(s)}, Y^{(s)})$ is a subset of $(X, Y)$. It follows that

$$\text{if } Var[X] > Var[X^{(s)}] \text{ then } \rho_{XY} > \rho_{XY^{(s)}} > 0, \text{ where } s \subsetneq S \tag{9}$$

Since $G(.)$ in Eq.(7) is a monotonic function, plugging Eq.(7) into Eq.(9) gives

$$G(\rho_{XY}) > G(\rho_{XY^{(s)}}) > 0 \Rightarrow \tau_{XY} > \tau_{XY^{(s)}} > 0 \text{ given that } Var[X] > Var[X^{(s)}] \text{ where } s \subsetneq S \tag{10}$$

**Lemma 3:** The overall concordance can be written as the weighted mean of the concordances of non-overlapping sets of pairs.

Let $\Phi(\langle i,j \rangle | X, Y) = \Phi(i,j)$ be a pairwise concordance operator, which returns 1 if the observation pair $\langle i,j \rangle$ is concordant on $(X, Y)$, and 0 otherwise:

$$\Phi(i,j) = \begin{cases} 1 & \text{if } sgn(x_i - x_j) = sgn(y_i - y_j) \\ 0 & \text{otherwise} \end{cases} \tag{11}$$

The overall concordance $d = d_{YX}$ between $Y$ and $X$ can be formulated in terms of the operator $\Phi$:

$$d = \frac{1}{|\pi|} \sum_{\langle i,j \rangle \in \pi} \Phi(i,j) = \mathbb{E}[\, \Phi(i,j) \mid \langle i,j \rangle \in \pi \,] = \mathbb{E}[\, \Phi_\pi \,] \tag{12}$$

where $\pi$ is the set of all pairs. This means that concordance can be expressed as the expected value of the pairwise ranking operator over the set of all pairs. Considering $\pi = \pi_1 \cup \pi_b \cup \pi_2$, we can write the overall concordance as:

$$d = \mathbb{E}[\Phi_\pi] = \frac{|\pi_1|}{|\pi|}\mathbb{E}[\Phi_{\pi_1}] + \frac{|\pi_b|}{|\pi|}\mathbb{E}[\Phi_{\pi_b}] + \frac{|\pi_2|}{|\pi|}\mathbb{E}[\Phi_{\pi_2}] \tag{13}$$

Using Eq.(12), this can be written as:

$$d = \frac{|\pi_1|}{|\pi|} d_1 + \frac{|\pi_b|}{|\pi|} d_b + \frac{|\pi_2|}{|\pi|} d_2 \tag{14}$$

**Lemma 4:** Given a bivariate distribution of random variables $X$ and $Y$, with $Y$ being a continuous random variable with no ties, Kendall's rank correlation coefficients $\tau$, $\tau^a$, and concordance $d$ are monotonically related due to the following equality [13]:

$$\tau_{YX} = \frac{\tau^a_{YX}}{\tau^a_{YY}} = 2d_{YX} - 1 \tag{15}$$

Note that $\tau^a$ is a variant of Kendall's $\tau$ which counts ties as discordant [36].

##### Main Theorem

**Theorem:** For $d > 0.5$, if an LOD effect exists $\Rightarrow d_b > d > d_1$

**Proof:** Assume that there exist monotonic transformation functions $t_x$ and $t_y$, such that $t_x(X) = \tilde{X}, t_y(Y) = \tilde{Y}$, with $\tilde{X}\tilde{Y} = (\tilde{X}, \tilde{Y})$ being a bivariate normal distribution. Given the monotonicity of the transformation, $(X, Y)$ and $(\tilde{X}, \tilde{Y})$ are rank-invariant, and therefore any rank-based statistic on the transformed data will produce results equivalent to the ones on the original data. For example, if we calculated Kendall's $\tau$, this would mean $\tau_{YX} = \tau_{t_x(Y)t_y(X)} = \tau_{\tilde{Y}\tilde{X}}$.

Now let $y_{th}$ be the LOD threshold on $Y$, and $s_2$ be the set of all observations with $Y$ value below the LOD: $s_2 = \{i \in S \mid y_i < y_{th}\}$, such that $min(Y) \leq \{y \in Y^{(s_2)}\} < y_{th} < \{y \in Y^{(s_1)}\} \leq max(Y)$. In the transformed space, this becomes

$$\begin{aligned} min(\tilde{Y}) \leq \tilde{y} < \tilde{y}_{th} \quad \forall \tilde{y} \in \tilde{Y}^{(s_2)} \\ max(\tilde{Y}) \geq \tilde{y} > \tilde{y}_{th} \quad \forall \tilde{y} \in \tilde{Y}^{(s_1)} \end{aligned} \tag{16}$$

In an ordered vector of continuous values, the variance of the overall vector is always larger than that of any subsets of of adjacent elements. For example, given an ordered vector $v = \{v_1, v_2, ...v_n\}$ with elements $\{v_i \in R \mid v_{i-1} < v_i,\ i = 1, 2, ...n\}$, and an ordered subvector $k \subsetneq v$, say $k = \{v_1, v_2, v_3\}$, then Var$[v] >$ Var$[k]$. Applying this observation to our subsets, we deduce that Var$[\tilde{Y}] > \{$Var$[\tilde{Y}^{(s_1)}]$, Var$[\tilde{Y}^{(s_2)}]\}$. Substituting this inequality into Eq.(10) leads to $\tau_{\tilde{Y}\tilde{X}} > \tau_{\tilde{Y}\tilde{X}^{(s_2)}}, \tau_{\tilde{Y}\tilde{X}^{(s_1)}}$, namely that Kendall's correlation between $\tilde{X}$ and $\tilde{Y}$ on the whole set of observations $S$ must be larger than the Kendall's correlation computed on either subset $s_1$, $s_2$. Since $\tilde{Y}$ is a continuous random variable, we can write the concordance between $\tilde{Y}$ and $\tilde{X}$ as $d = (\tau + 1)/2$ (Lemma 4). This, taken together with the previous $\tau$ inequality, leads to $d > \{d_1, d_2\}$. From Lemma 3 we know that the overall concordance $d$ is a weighted average of $d_1, d_2$, and $d_b$. Since $d > \{d_1, d_2\}$, it must follow that $d_b > d$, and therefore

$$d_b > d > d_1, d_2 \tag{17}$$

This means that, if a limit of detection (LOD) exists, the concordance pertaining to the bridge pairs $d_b$ must be greater than the concordance pertaining to data above the LOD, i.e., the observed data points, $d_1$. Thus,

$$\text{if LOD} \Rightarrow d_b > d > d_1 \tag{18}$$

□

**Remark:** The true concordance is a weighted mean of $d_1, d_b, d_2$ as shown in Eq.(14). Considering $d > d_2$ from Eq.(17), once $d_2$ is dropped from the weighted average as in the *rox* core model Eq.(1), the concordance will increase. Thus, without further adjustment, the estimated concordance will be higher than the true value and thus be overestimated.

##### Supplementary Text 3: Choice of weight parameter $p$ for debiasing

Here, we will derive the debiasing formulation of $rox$ as presented in Eq.(3). First, we will derive an approximation for the probability of a single pair being concordant. Then using this approximation, we will derive the debiasing weight for the overall concordance of $rox$ as presented in the Eq.(3).

##### An approximation for the probability of a pair of observations being concordant

Let $Y$ and $X$ be two positively correlated random variables with $d_{YX} > 0.5$. In this setting, we will first obtain an approximation for the probability of a pair being concordant on $(X, Y)$ given the ranks of each data point in $Y$: $P(\Phi(i,j) = 1|r_i, r_j) = P(\Phi_{ij} = 1)$, where $\Phi(i,j)$ is the pairwise ranking operator defined in Lemma 3, Eq.(11), and $r_i = \text{rank}(y_i)/\max(\text{rank}(Y))$ is the scaled ranking of data point $y_i$. Note that, given the ranks on $Y$, we derive an approximation for the probability of a pair being concordant, not for overall concordance. Then using this probability, we will derive the debiasing weight of $rox$ as presented in the Eq.(3).

**Theorem:** Given two random variables $X, Y$ with $d_{YX} > 0.5$, there exists a linear function $f(x) = mx + k, m \in \mathbb{R}^+$ $k \in \mathbb{R}$ such that

$$P(\Phi_{ij} = 1|r_i, r_j) \approx f(r_i - r_j)$$

**Proof:** Let $Y^r = \text{rank}(Y)/\max(\text{rank}(Y))$ be the scaled rank-transformed version of $Y$. Assume that a monotonic transformation function $t_x$ exists such that $X^t = t_x(X)$, and $X^t = \beta Y^r + \epsilon$, $\beta > 0$. Define a random variable $\zeta = \epsilon_1 - \epsilon_2$, where $\epsilon_1$ and $\epsilon_2$ have the same distribution of $\epsilon$. Hence, $\zeta$ follows a symmetric distribution centered at 0 with $Var[\zeta] = 2Var[\epsilon]$. Let $F_\zeta$ be the cumulative distribution function (CDF) of $\zeta$.

Given a pair of observations $\langle i, j\rangle$, a pair of ranks $\{r_i > r_j | r_i, r_j \in Y^r\}$, and a pair of values $\{x_i^t, x_j^t \in X^t\}$, then the probability of $\langle i, j\rangle$ being concordant is:

$$\begin{aligned} P(\Phi_{ij} = 1) &= P\left[\left(x_i^t - x_j^t\right) > 0\right] = P((\beta r_i + \epsilon_i - \beta r_j - \epsilon_j) > 0) \\ &= P(\beta(r_i - r_j) > (\epsilon_i - \epsilon_j)) = P\left(\beta(r_i - r_j) > \zeta_{ij}\right) \\ &= F_\zeta\left[\beta(r_i - r_j)\right] \approx m\,(r_i - r_j) + k \ \text{ with } \ m > 0 \end{aligned} \tag{19}$$

Explanation of the approximation in the last line of the equation: All CDFs (like $F_\zeta$) are monotonically increasing functions bounded by $[0, 1]$. Any continuous monotonic function on a closed interval can be approximated with a simple linear function. The goodness of such approximation depends on the shape of CDF to be approximated. $\zeta$ follows a symmetric distribution with $Var[\zeta] = 2Var[\epsilon]$. For a symmetric distribution, a higher variance translates into a less "steep" sigmoid of the corresponding CDF. Notably, the less steep the sigmoid curve, the better it can be approximated by a linear function.

□

**Remark:** The assumption that a monotonic transformation function $t_x(X) = X^t$ exists such that $X^t = \beta Y^r + \epsilon$, with $\beta > 0$, is a mild assumption, because a simple rank transformation would always satisfy this condition. Let $t_x$ be the rank transformation such that $X^t = t_x(X)/\max(t_x(X))$ is the scaled rank transform of $X$, then $\text{rank}(X) = \beta\,\text{rank}(Y) + \epsilon$, which is related to Spearman correlation $\rho^S_{YX}$.

Since $d_{YX} > 0.5$ by design, and $\tau_{YX} = 2d_{YX} - 1$ (Lemma 3), and $\tau_{YX} > \rho^S_{YX}$ [37], we can conclude that $\rho^S_{YX} > \tau_{YX} > 0$, and hence $\beta > 0$.

##### Debiasing weight

In Eq.(19), we obtained an approximation for the probability of a pair being concordant as $P(\Phi_{i,j} = 1) = f(r_i - r_j)$, which we now use to formulate the overall concordance $d$. As shown in Lemma 3, Eq.(12), concordance can be expressed as the expected value of correct pairwise rankings: $d = \mathbb{E}[\Phi_\pi]$, where $\pi$ is the set of all pairs and $\Phi$ is the pairwise ranking operator defined therein.

$$\begin{aligned} d = \mathbb{E}[\Phi_\pi] &= \frac{1}{|\pi|} \sum_{\langle i,j\rangle \in \pi} \mathbb{E}_{\Phi_{ij}} = \frac{1}{|\pi|} \sum_{\langle i,j\rangle \in \pi} P(\Phi_{ij} = 1) \\ &\approx \frac{1}{|\pi|} \sum_{\langle i,j\rangle \in \pi} f(r_i - r_j) = f\left(\frac{1}{|\pi|} \sum_{\langle i,j\rangle \in \pi} (r_i - r_j)\right) \text{ ( because of linearity)} \\ d &\approx f(\Delta), \end{aligned} \tag{20}$$

where $\Delta = \frac{1}{|\pi|} \sum_{\langle i,j\rangle \in \pi} (r_i - r_j)$ is the mean of scaled ranking difference among pairs in $\pi$.

As shown in Lemma 3, the overall concordance can be expressed as the weighted mean of the concordances of non-overlapping sets of pairs. If the set of samples $S$ was divided into two non-overlapping groups $s_1$ and $s_2$ such that $S = s_1 \cup s_2$ (see Supplementary Figure 7), we can divide all pairs $\pi$ into three sets of pairs: $\pi_1$, which includes the pairs within first group, $\pi_2$, which includes the pairs within second group, and $\pi_b$, which includes the pairs between two groups, i.e., the bridge pairs. We thus have $\pi = \pi_1 \cup \pi_b \cup \pi_2$, and hence $d = \frac{|\pi_1|}{|\pi|} d_1 + \frac{|\pi_b|}{|\pi|} d_b + \frac{|\pi_2|}{|\pi|} d_2 = \frac{|\pi_1|d_1 + |\pi_2|d_2 + |\pi_b|d_b}{|\pi_1| + |\pi_2| + |\pi_b|}$, where $d, d_1, d_2, d_b$ are the concordances pertaining to $S, s_1, s_2$, and the bridge pairs, respectively.

The goal is to find a weight $w$ in the absence of $d_2$ by weighting samples of $s_2$ such that the concordance computed on the observable pairs is equal to the true concordance, i.e., $\hat{d}(w) = d$. Let $n_1$ and $n_2$ be the sample sizes of $s_1$ and $s_2$, respectively, and $n = n_1 + n_2$ be the overall sample size,

$$d = \frac{|\pi_1|d_1 + |\pi_2|d_2 + |\pi_b|d_b}{|\pi_1| + |\pi_2| + |\pi_b|}, \; \hat{d}(w) = \frac{|\pi_1|d_1 + w|\pi_b|d_b}{|\pi_1| + w|\pi_b|} \tag{21}$$

By requiring that $\hat{d}(w) = d$, we get

$$\begin{aligned} d = \hat{d}(w) &\Rightarrow \frac{|\pi_1|d_1 + |\pi_2|d_2 + |\pi_b|d_b}{|\pi_1| + |\pi_2| + |\pi_b|} = \frac{|\pi_1|d_1 + w|\pi_b|d_b}{|\pi_1| + w|\pi_b|} \\ &\Rightarrow (|\pi_1|d_1 + |\pi_2|d_2 + |\pi_b|d_b)(|\pi_1| + w|\pi_b|) = (|\pi_1|d_1 + w|\pi_b|d_b)|\pi| \\ &\Rightarrow w = \frac{|\pi_1|\left(|\pi_1|d_1 + |\pi_2|d_2 + |\pi_b|d_b\right) - |\pi||\pi_1|d_1}{|\pi||\pi_b|d_b - |\pi_b|(|\pi_1|d_1 + |\pi_2|d_2 + |\pi_b|d_b)} \end{aligned} \tag{22}$$

In Eq.(20), we derived an approximation for the concordance $d = f(\Delta)$, where $\Delta$ is the mean pairwise ranking difference on $X$. Let us first calculate $\Delta$ for $\pi_1$, $\pi_2$, and $\pi_b$:

$$\begin{aligned} \Delta_1 &= \frac{1}{|\pi_1|} \sum_{\langle i,j\rangle \in \pi_1} (r_i - r_j) = \frac{1}{n_1(n_1-1)/2} \sum_{j=1}^{n_1-1} \sum_{i=j+1}^{n_1} \frac{1}{n}(i-j) = \frac{n_1+1}{3n} \\ \Delta_2 &= \frac{1}{|\pi_2|} \sum_{\langle i,j\rangle \in \pi_2} (r_i - r_j) = \frac{1}{n_2(n_2-1)/2} \sum_{j=1}^{n_2-1} \sum_{i=j+1}^{n_2} \frac{1}{n}(i-j) = \frac{n_2+1}{3n} \\ \Delta_b &= \frac{1}{|\pi_b|} \sum_{\langle i,j\rangle \in \pi_b} (r_i - r_j) = \frac{1}{n_1 n_2} \sum_{j=1}^{n_2} \sum_{i=j+1}^{n_1} \frac{1}{n}(i-j) = \frac{n_1+n_2}{2n} = \frac{n}{2n} \end{aligned} \tag{23}$$

Using Eq.(20), these individual concordances can be written based on a linear approximation. Considering the $\Delta_1, \Delta_2$, and $\Delta_b$ we obtained, we can write $d_1, d_2, d_b$ as follows:

$$\begin{aligned} d_1 \approx f(\Delta_1) &\Rightarrow d_1 \approx f\left(\frac{n_1+1}{3n}\right) \\ d_2 \approx f(\Delta_2) &\Rightarrow d_2 \approx f\left(\frac{n_2+1}{3n}\right) \\ d_b \approx f(\Delta_b) &\Rightarrow d_b \approx f\left(\frac{1}{2}\right) \end{aligned} \tag{24}$$

After this, all concordances formulated in Eq.(24) are only based on $f$ and sample sizes. Plugging these approximations into Eq.(22), and using the fact that $|\pi| = n(n-1)/2$, $|\pi_1| = n_1(n_1-1)/2$, $|\pi_2| = n_2(n_2-1)/2$, and $|\pi_b| = n_1 n_2$, we get

$$
\begin{aligned}
w &\approx \frac{|\pi_1||\pi_1|f\left(\frac{n_1+1}{3}\right) + |\pi_1||\pi_2|f\left(\frac{n_2+1}{3}\right) + |\pi_1||\pi_b|f\left(\frac{n}{2}\right) - |\pi||\pi_1|f\left(\frac{n_1+1}{3}\right)}{|\pi||\pi_b|f\left(\frac{n}{2}\right) - |\pi_b||\pi_1|f\left(\frac{n_1+1}{3}\right) - |\pi_b||\pi_2|f\left(\frac{n_2+1}{3}\right) - |\pi_b||\pi_b|f\left(\frac{n}{2}\right)} \\
&= \frac{|\pi_1||\pi_1|\frac{n_1+1}{3n} + |\pi_1||\pi_2|\frac{n_2+1}{3n} + |\pi_1||\pi_b|\frac{1}{2} - |\pi||\pi_1|\frac{n_1+1}{3n}}{|\pi||\pi_b|\frac{1}{2} - |\pi_b||\pi_1|\frac{n_1+1}{3n} - |\pi_b||\pi_2|\frac{n_2+1}{3n} - |\pi_b||\pi_b|\frac{1}{2}} \quad \text{($m$'s and $k$'s in $f(x) = mx + k$ are cancelled out)} \\
&= \frac{\left(\frac{n_1(n_1-1)}{2}\right)^2 \frac{n_1+1}{3} + \frac{n_1(n_1-1)}{2}\frac{n_2(n_2-1)}{2}\frac{n_2+1}{3} + \frac{n_1(n_1-1)}{2} n_1 n_2 \frac{n}{2} - \frac{n(n-1)}{2}\frac{n_1(n_1-1)}{2}\frac{n_1+1}{3}}{\frac{n(n-1)}{2} n_1 n_2 \frac{n}{2} - n_1 n_2 \frac{n_1(n_1-1)}{2}\frac{n_1+1}{3} - n_1 n_2 \frac{n_2(n_2-1)}{2}\frac{n_2+1}{3} - (n_1 n_2)^2 \frac{n}{2}} \\
&= \frac{n_1 n_2 (n_1-1)\left(n_1^2 + 2n_1 n_2 - n_1 + n_2^2 - n_2\right)}{n_1 n_2 \left(n_1^3 + 3n_1^2 n_2 - 3n_1^2 + 3n_1 n_2^2 - 6n_1 n_2 + 2n_1 + n_2^3 - 3n_2^2 + 2n_2\right)} \\
&= \frac{n_1 n_2 (n_1-1)(n_1+n_2)(n_1+n_2-1)}{n_1 n_2 (n_1+n_2-2)(n_1+n_2)(n_1+n_2-1)} = \frac{n_1-1}{n_1+n_2-2} = \frac{n_1-1}{n-2}
\end{aligned} \tag{25}
$$

$$
\Rightarrow w = \frac{n_1-1}{n-2} \approx \frac{n_1}{n} = 1 - \frac{n_2}{n} \tag{26}
$$

□

#### Simulation

To verify that the proposed approximated weight from Eq.(26) performs well compared to the optimal weight from Eq.(22), we performed an extensive simulation with various ground truths and missingness percentages.

Let $X, Y$ be random variables, and $d = d_{YX}$ the concordance between them. We simulated both $X$ and $Y$ as normally distributed $Y = X + \alpha\epsilon$ where $Y \sim \mathcal{N}(\mu_y, \sigma_y^2)$, $\epsilon \sim \mathcal{N}(\mu_\epsilon, \sigma_\epsilon^2)$, and $\alpha \in \mathbb{R}^+$. We tweaked the noise factor $\alpha$ in order to obtain variables with overall concordance $d = \{0.55, 0.60, 0.65, 0.70, 0.75, 0.80, 0.85\}$. To simulate the ground truth robustly, we generated $n = 10^4$ observations. In this simulated scenarios, we calculated the optimal weight based on Eq.(22) such that $d(w_{opt}) = d$. We then introduced varying proportions of missing values, for which we recorded the optimal weight $w_{opt}$, as well as the proposed approximate weight $w = 1 - n_2/n$. By comparing then $w$ with $w_{opt}$ we observed that the proposed weight was remarkably close to the optimal weight (see Supplementary Figure 8).

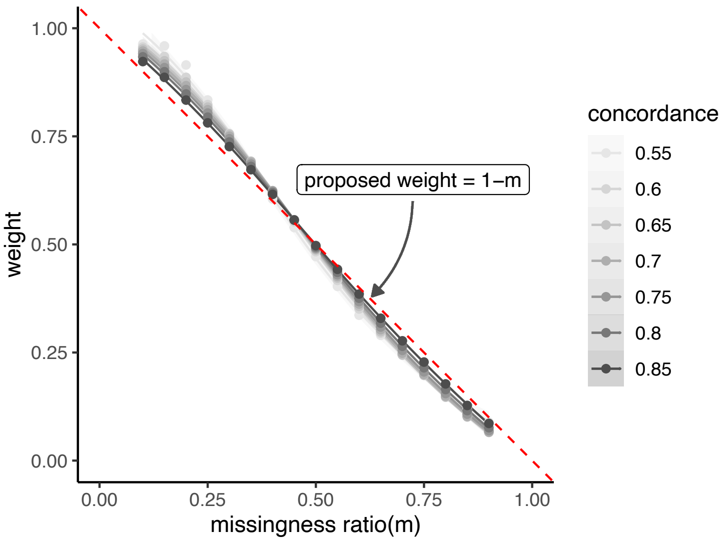

Supplementary Figure 8: For each $d$, missingness was introduced at a rate of $10\%, 15\%, \ldots, 90\%$ of all samples simulated. Solid lines represent the optimal weight $w_{opt}$ for each $d$, and dashed red line represent the proposed approximate weight $w$, which is only dependent on the missingness percentage.

**Supplementary Figure 1: Simulation framework and small sample size results**

For small sample sizes in the simulation framework, we again compared $rox$ to complete case analysis (CCA) and minimum imputation (min-imp) with regular concordance analysis. $Y$ and $X$ were simulated 1,000 times for $n = 100$ with concordance values $d = 0.6, 0.7, 0.8$. Missing values were then introduced in $Y$ with varying percentages and degree of LOD effect. For both strict LOD with differing missingness percentages (panel A) and probabilistic LOD with varying LOD effects (panel B), $rox$ demonstrated overall superior performance compared to min-imp and CCA.

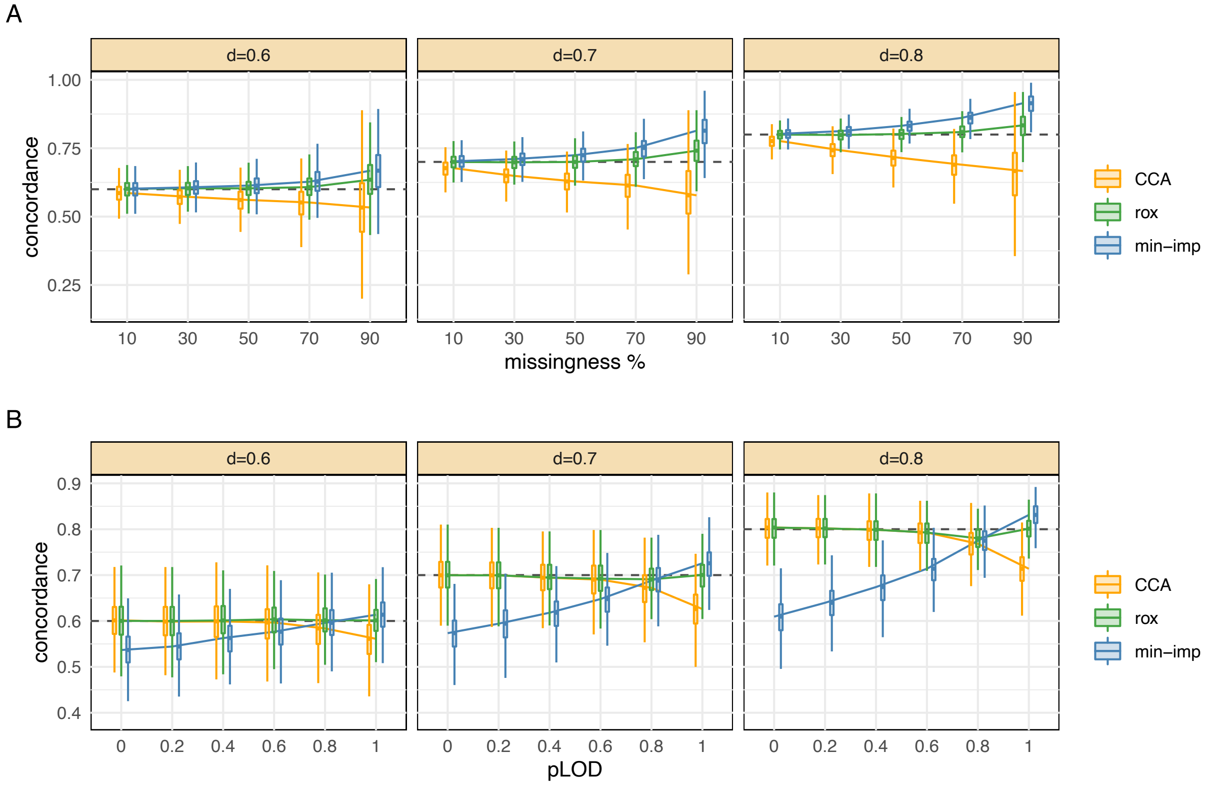

**Supplementary Figure 2: Simulation in multivariable setting**

Data was simulated as i.i.d. with sample size $n = 10,000$; see METHODS for details on the simulation approach. Strict LOD was simulated with varying missingness percentages and probabilistic LOD was simulated with 50% missingness with varying LOD effect strengths. In this framework, the multivariable extension of $rox$ was compared with ordinary linear regression after minimum imputation (lm+min) and ordinary linear regression with complete case analysis (lm+CCA). Performance was assessed by the ability of the models to recover the true regression coefficients. Note that the true concordance and $\beta$ coefficent were determined by running regression analysis on the simulated data without missing values.

Under strict LOD, the $rox$ model performed better than lm+min-imp and lm+CCA in recovering the regression coefficients and overall model fit. Under probabilistic-LOD, $rox$ and lm+CCA performed similarly in terms of recovering the true regression coefficients when the LOD effect was not prominent (i.e., for $pLOD < 0.7$). The estimates of regression $\beta$ coefficients after minimum imputation were biased for all simulation parameters. Under a prominent LOD effect, $rox$ generally performed better than both other approaches.

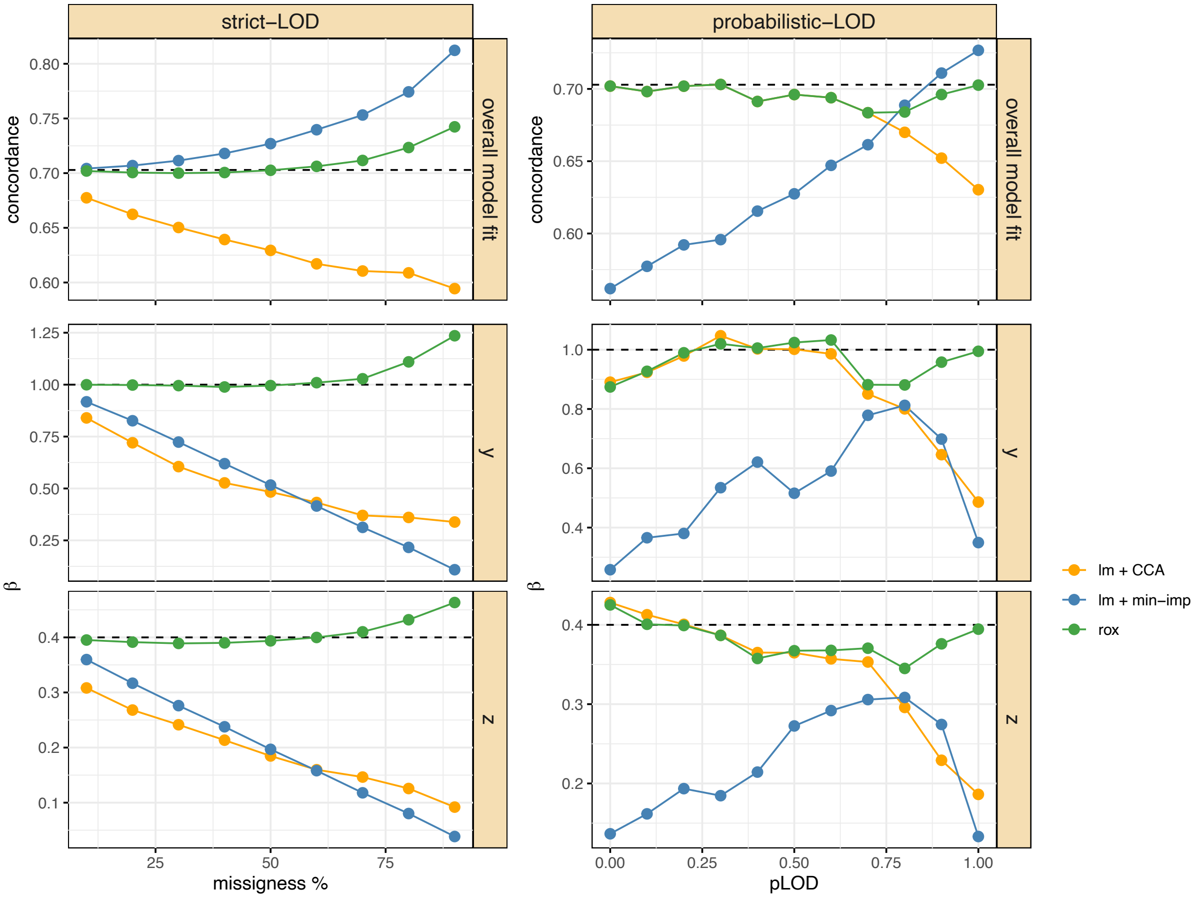

**Supplementary Figure 3: Estimating degree of LOD effect in the QMDiab data**

A total of 292 samples in the QMdiab study were profiled using two different metabolomics platforms, which cover many of the same metabolites. We here provide evidence for the assumption of an LOD effect in the data. Since each platform measured the same metabolite on the same samples but with different missingness, insights into whether missing values of a metabolite are due to LOD can be gained from the corresponding non-missing measurements of the other platform. Intuitively, if a value is missing on one platform and measured on the other platform, those values are expected to be on the low end of the distribution.

We estimated the LOD effect based on an ROC-AUC analysis, where classes corresponds to missing yes vs. no on one platform, and scores correspond to the measurement values on the other platform. For each metabolite measured on both platforms, we considered the measurement from the platform with the lower number of missing values as our reference, and used the other platform values for comparison. Observations with missing values in the reference platform were discarded. Formally, the AUC is formulated as $AUC_{LOD} = P(x_i^{ref} < x_j^{ref} | x_i^{pl} = NA, x_j^{pl} \neq NA)$ where $x^{ref}$ represents the measured values from the reference platform and $x^{pl}$ represents the corresponding observations in the other platform. In other words, a potential LOD effect was estimated as the probability of a value being missing in one dataset given that its measured value in the other dataset is lower than all measured values.

An $AUC = 1$ for a given metabolite indicates that for all missing values in the reference sample, the correpoonding values in the comparison sample were lower than all other measurements. This indicates a strict LOD-based missingness pattern. In contrast, an $AUC = 0.5$ indicates no overall separation of missing and measured values between reference and comparison sample, hence indicating a "missing-at-random" (MAR) pattern. The figure below illustrates the distribution of AUCs for all metabolites measured on both platforms, showing that most metabolites display a relatively strict LOD missingness pattern.

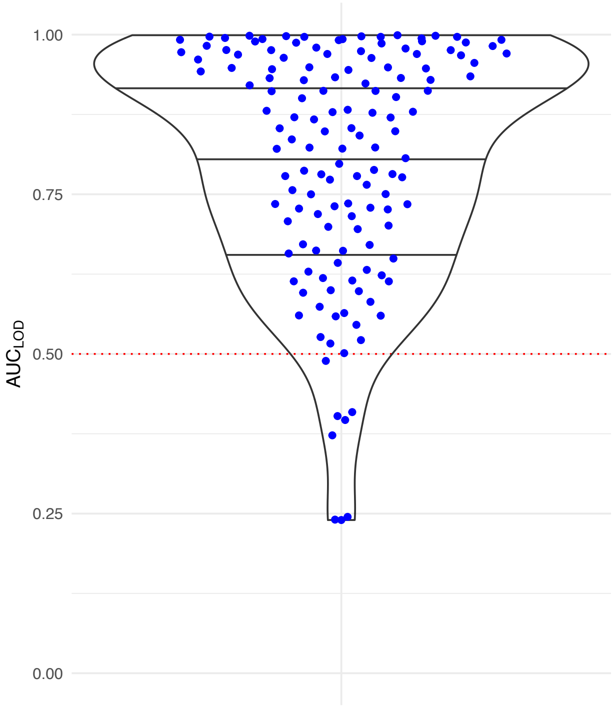

#### Supplementary Figure 4: Multi-Platform Validation - detailed results

This analysis extends the fully quantified (FQ) vs. partially missing (PM) ground truth analysis outlined in the main manuscript. (A) is the same panel as in the main manuscript, Figure 5B, showing systematic results for all four outcomes and varying levels of missingness in the PM metabolites. The x-axis indicates the minimum fraction of missing values per metabolite. The y-axis shows the Pearson correlation between the estimates across the two platforms. Panel (B): Detailed results across all outcomes where the partially missing metabolite has at least 20% missingness. Points distributed along the diagonal indicate consistent estimations between the two platforms. The gray line indicates the x=y axis, while the colored line indicates a linear fit over the points of comparison. Across all missingness percentages, $rox$ was more consistent compared to knn imputation and similarly or more consistent compared to minimum imputation.

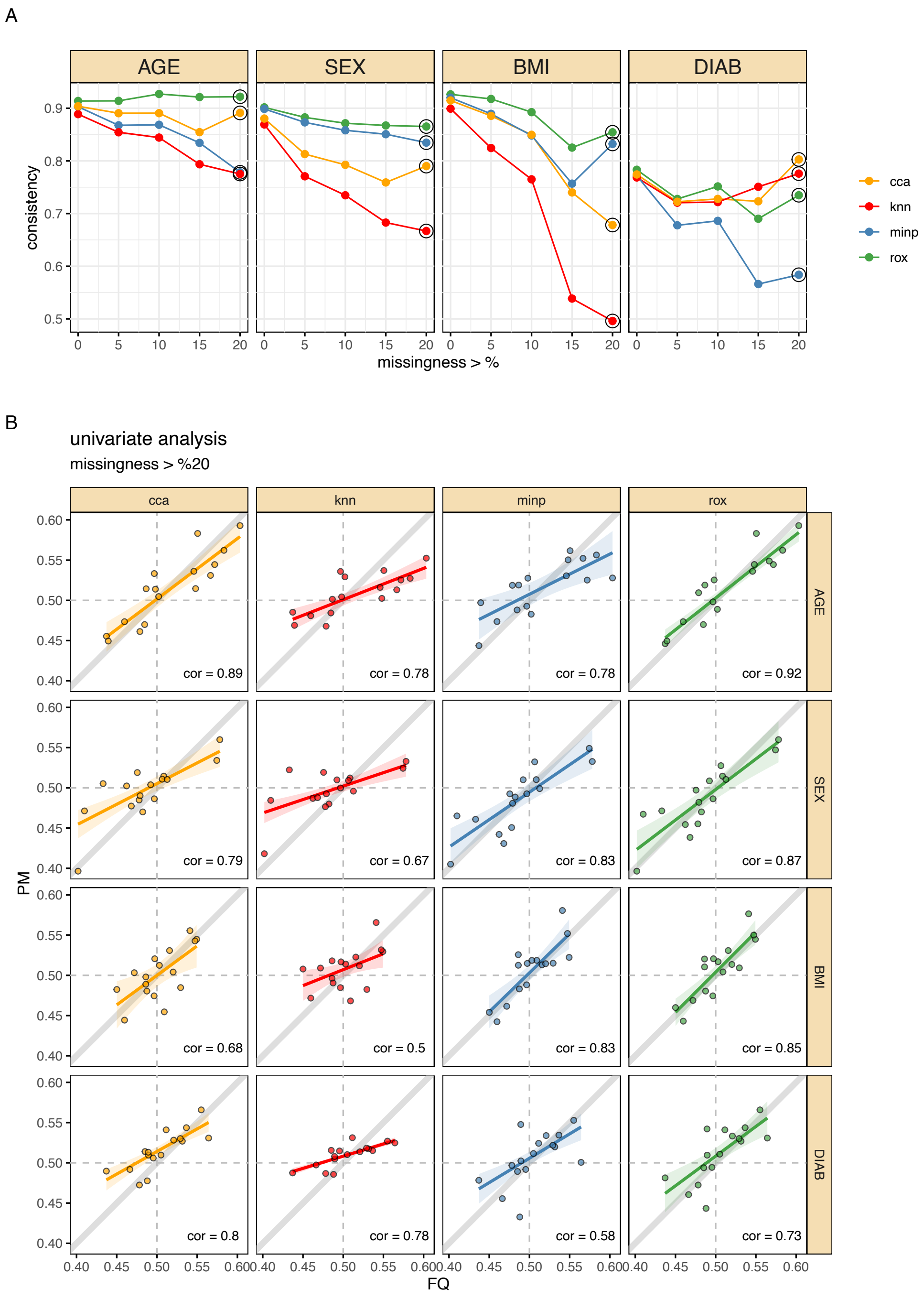

**Supplementary Figure 5: Multi-Platform Validation - multivariable model**

Here we extended the fully quantified (FQ) vs. partially missing (PM) ground truth analysis outlined in the main manuscript to the multivariable modeling case. To this end, we tested all four outcomes in a single model, i.e., $metabolite \sim age + sex + bmi + diabetes$. For each metabolite, regression coefficients were calculated for two platforms independently. We compared $rox$ to three other common approaches: (1) Linear regression after dropping missing values (CCA+lm), (2) linear regression after knn imputation (knn+lm), and (3) linear regression after minimum imputation (minp+lm). The consistency of results between the regression coefficients from the FQ and the PM analysis were computed with Pearson correlation. Independent of missingness percentage, $rox$ displayed similar or more consistent coefficient estimation compared to its competitors for all variables (panel A). Detailed consistency calculation results for BMI as an example are shown in panel B.

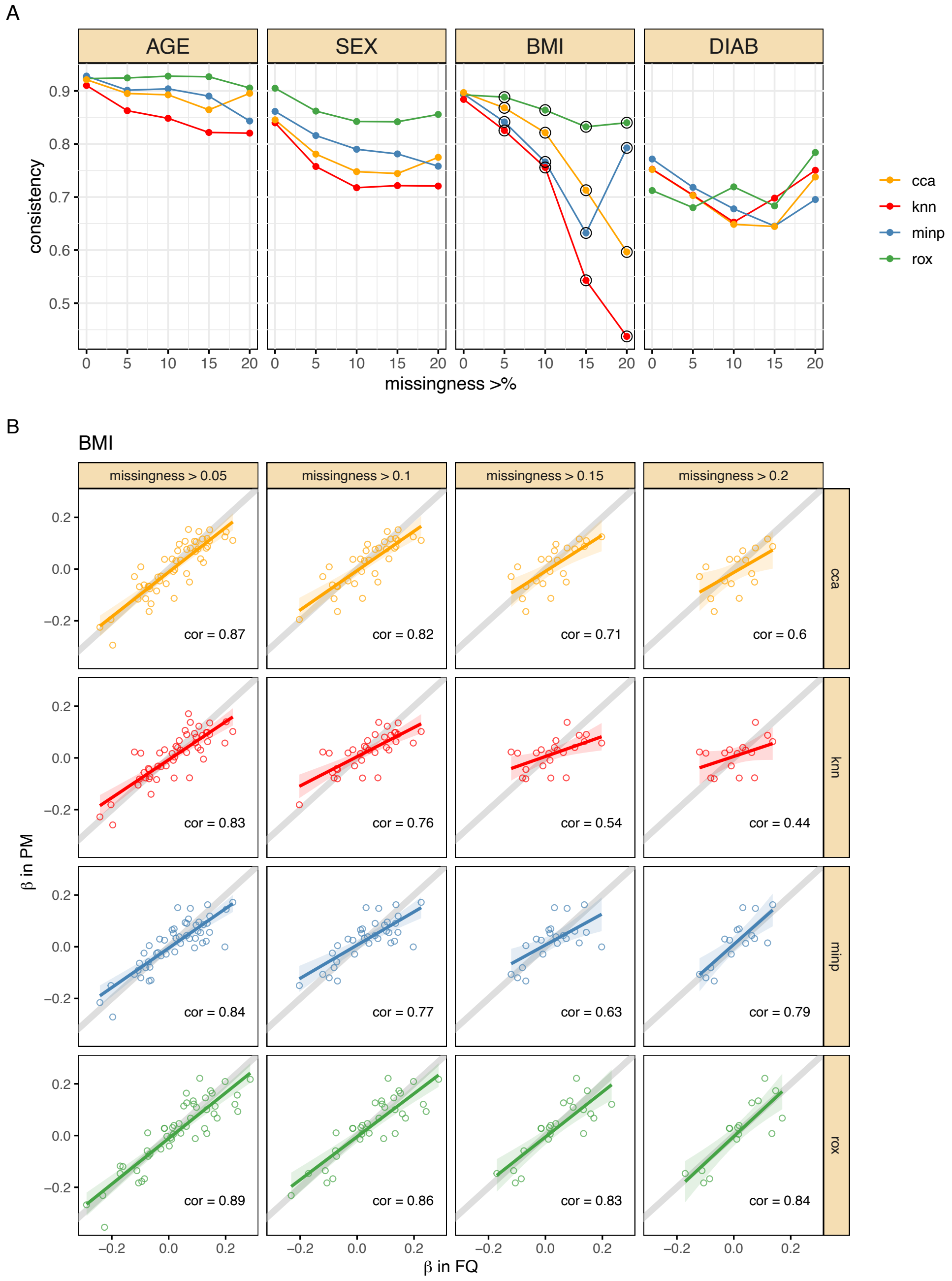

**Supplementary Figure 6: Comparable pairs with left-censoring**

Illustration of observed ($s_1$) and unobserved data ($s_2$) points, as well as pairs between measured data points ($\pi_1$), unmeasured data points ($\pi_2$) and the "bridge" pairs in between ($\pi_b$).

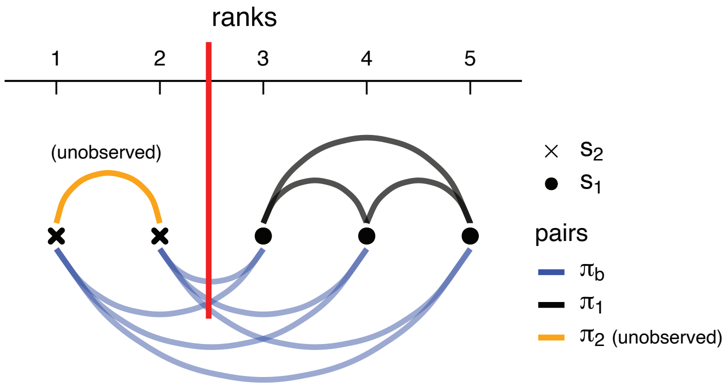
